# Biosynthesis of Kaitocephalin: A Neuroprotective Natural Product Featuring a Peptide-Like yet Non-Peptidic Scaffold

**DOI:** 10.1101/2025.10.18.683206

**Authors:** Yukari Maeno, Taro Shiraishi, Naoya Saito, Jun-ichi Maruyama, Kazuo Shin-ya, Tomohisa Kuzuyama

## Abstract

Kaitocephalin (KCP, **1**) is a neuroprotective natural product that acts as an antagonist of ionotropic glutamate receptors, making it a highly promising lead for drug discovery. It possesses a unique scaffold composed of three amino acids connected via C–C bonds, which appears peptide-like but is formed without peptide bonds. In this study, we identified the KCP biosynthetic gene cluster (*kpb* cluster) in the producing fungus *Eupenicillium shearii* through integrated genomic and transcriptomic analyses. LC-MS/MS profiling and chemical derivatization of *E. shearii* extracts led to the discovery of four novel pathway-related metabolites (**2**–**5**). *In vitro* enzymatic assays with 2(*S*)-dechlorokaito lactate (**4**) as a substrate enabled functional characterization of KpbI, KpbM, and KpbB involved in KCP formation. Among them, the dioxygenase KpbI was found to catalyze an unprecedented two-step oxidation to form the D-serine moiety. In addition, isotope tracing experiments provided new insights into the origin of the L-proline moiety. These findings establish a foundation for future studies aimed at elucidating the complete biosynthetic mechanism of KCP.

**Table of Contents graphical abstract:** 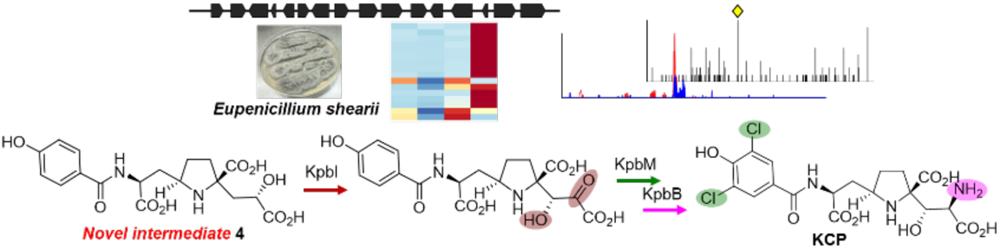

## Introduction

Ionotropic glutamate receptors (iGluRs) are ligand-gated ion channels and mediate excitatory neurotransmission in the mammalian central nervous system. Excessive activation of iGluRs by glutamate can lead to neuronal injury or death, a process known as excitotoxicity.^1,2^ Kaitocephalin (KCP, **1**, Figure 1) was isolated from the filamentous fungus *Eupenicillium shearii* PF1191 through screening based on its neuroprotective activity against excitotoxicity.^3^ Biochemical and electrophysiological studies have demonstrated that KCP functions as an antagonist of iGluRs, exhibiting high subtype selectivity for *N*-methyl-D-aspartate (NMDA) receptors over the other two subtypes AMPA and kainate receptors.^4^ NMDA receptors are ubiquitously dispersed throughout the central nervous system, play crucial roles in brain development and function, and are the targets of clinically relevant drugs for the treatment of mild cognitive impairment, schizophrenia, depression, and epilepsy.^5^ Memantine, an NMDA receptor antagonist, has been shown to attenuate the progression of symptoms in patients with moderate to severe Alzheimer’s disease,^6^ and it has been approved as a therapeutic agent for this condition.

**Figure 1.**
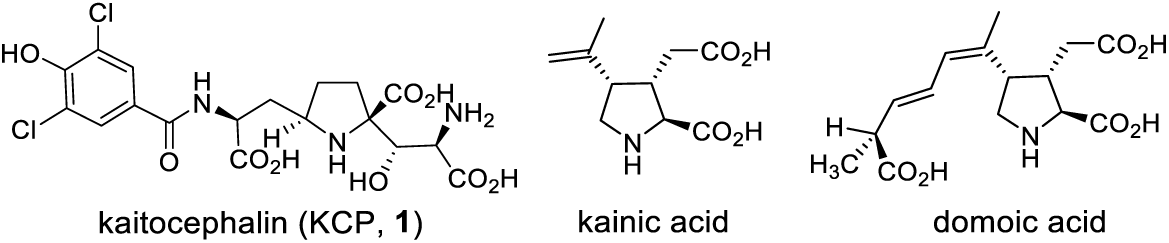
Natural product ligands for iGluRs.

Thus, KCP, which has remarkable bioactivity with promising pharmaceutical applications, has also drawn considerable interest from chemists because of its unique chemical structure. KCP comprises a three-amino-acid unit of L-alanine, L-proline, and D-serine, together with an amide-linked 3,5-dichloro-4-hydroxybenzoyl moiety. The three amino acids are connected not through peptide bonds but via C–C bonds, resulting in a scaffold that appears peptide-like at first glance but is in fact entirely non-peptidic. KCP has been recognized as an attractive and challenging target for chemical synthesis due to its unusual structure and multiple stereocenters. Over the years, extensive efforts have been devoted to the chemical synthesis of KCP ^7–12^ and its various non-natural analogues, which have served as the basis for structure–activity relationship (SAR) studies.^11, 13–15^

Despite such well-characterized pharmacology and intriguing structure, the biosynthesis of KCP has remained elusive for nearly three decades since its isolation. Experimental validation that these three amino acids are biosynthetically joined via such linkages has not been established. The decades-long gap in biosynthetic studies on KCP is attributable to both the poor productivity of its producer *E. shearii* and the distinctive structural features of KCP, which precluded homology-based genome mining with canonical NRPSs or other typical scaffold-forming enzymes. In recent years, there has been a growing demand for sustainable and efficient supply systems for bioactive molecules through biosynthetic systems. For example, biosynthetic studies of the natural products kainic acid^16,17^ and domoic acid^18,19^ (Figure 1), which are potent ligands of the subtype kainate receptors of iGluRs,^20–22^ have been developed into scalable production systems^17^ and preparation of analogues for SAR studies.^23^ Elucidation of the biosynthetic pathway of KCP may not only fascinate chemists as an elegant natural strategy for constructing complex molecules, but also contribute to the establishment of an efficient production system for KCP, which holds potential as a pharmaceutical agent.

In this study, to elucidate the biosynthesis of KCP, we adopted an integrated approach consisting of three main strategies: (i) genome-transcriptome analysis using the KCP-producing fungus *E. shearii*, (ii) comparative LC-MS/MS profiling of *E. shearii* and (iii) *in vitro* enzymatic assays. This multifaceted approach enabled the identification of the biosynthetic gene cluster (BGC) of KCP, the isolation of novel biosynthetically related compounds, and the functional characterization of three enzymes involved in the late-stage biosynthesis of KCP. During this process, we discovered an α-ketoglutarate-dependent dioxygenase that catalyzes an unprecedented two step oxidation. Ultimately, we elucidated the mechanism underlying the formation of the D-serine moiety and the timing of chlorination during KCP biosynthesis. In addition, stable-isotope tracing experiment provided new insights into the origin of the L-proline moiety of KCP.

## Results and Discussion

### Identification of *kpb* cluster by genome-transcriptome analysis

The genome of the KCP-producing strain *E. shearii* PF1191 was sequenced, yielding a draft assembly of approximately 34 Mb. Gene prediction was performed using a web tool Augustus^24^ with the *Aspergillus oryzae* species model, generating a comprehensive set of predicted protein-coding genes. Given that KCP bears two chlorine atoms on its aromatic ring, we hypothesized that a halogenase might be involved in its biosynthesis. To identify the halogenase, a local BLASTP search was conducted using the amino acid sequences of known fungal halogenases such as RadH^25^ and AclH, ^26^ as queries. This search revealed a single halogenase-encoding gene in the genome of *E. shearii*. Functional annotation of neighboring genes and manual inspection of the surrounding region suggested the presence of a 33 kb biosynthetic gene cluster (BGC) comprising 13 open reading frames (ORFs) (Figure 2A). However, gene prediction in eukaryotic genomes is often complicated by introns, and we considered it insufficient to define the BGC solely by the presence of a single halogenase. We therefore sought transcriptome data to confirm the ORFs and strengthen the evidence for the cluster’s involvement in KCP biosynthesis.

**Figure 2.**
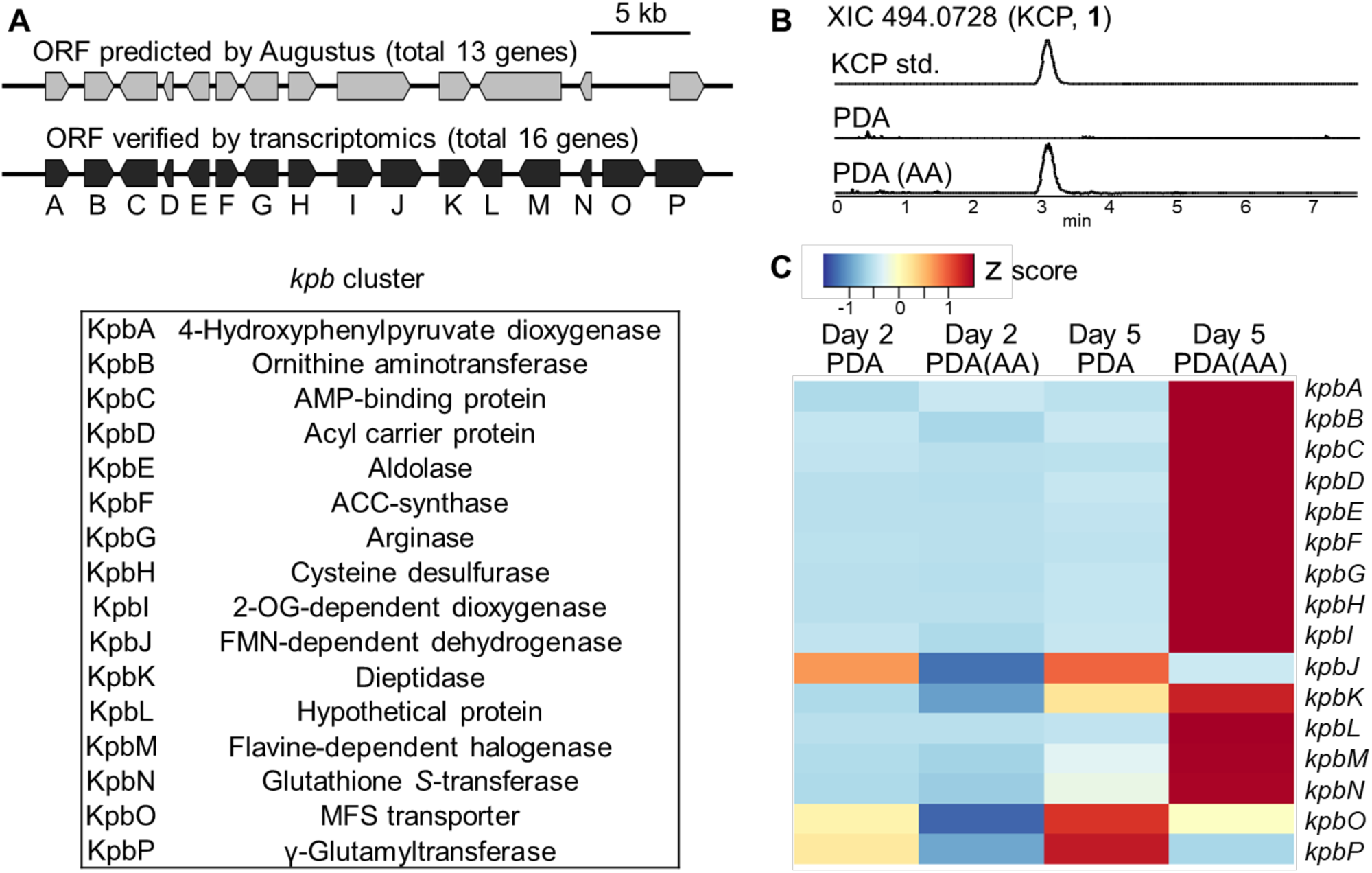
Identification of the *kpb* cluster by genome-transcriptome analysis. (A) ORFs within the *kpb* cluster predicted by Augustus (gray) and those verified by transcriptome analysis (black). The functional annotations of the *kpb* cluster genes are indicated. (B) Extracted ion chromatograms (XICs) at *m/z* 494.0728 corresponding to KCP (**1**). Shown are the KCP standard (top), extracts of *E. shearii* cultured for 9 days in PDA medium (middle), and extracts from PDA medium supplemented with adenine and L-arginine (bottom). “AA” denotes adenine and L-arginine supplementation. (C) Heatmap of gene expression levels. The three columns on the left represent KCP-nonproducing conditions, whereas the rightmost column represents the producing condition. “Day X” indicates the harvest day.

Because KCP is typically produced only in trace amounts,^3^ we first aimed to identify culture conditions that enhance its production and enable comparative transcriptome analysis. Extensive screening of the culture media revealed that KCP was reproducibly detected after 5 days of cultivation when both adenine and L-arginine were supplemented in potato dextrose agar (PDA) medium. In contrast, KCP was undetectable in the PDA medium lacking the supplements (Figure 2B) or when mycelia were harvested before Day 5, even in the presence of adenine and L-arginine. These results indicate that both supplementation and the cultivation duration are crucial for KCP production. We therefore conducted comparative transcriptome analysis using RNA extracted from mycelia grown under four conditions: with or without adenine and L-arginine supplementation and harvested on either Day 2 or Day 5. RNA-seq analysis revealed that the predicted cluster region actually contains 16 ORFs rather than 13 (Figure 2A). Moreover, most genes in the cluster were highly expressed only under the producing condition (Figure 2C). The results of the combined genome-transcriptome analysis strongly support the identification of the BGC responsible for KCP biosynthesis, which we designated as the *kpb* (kaitocephalin biosynthesis) cluster.

The functional annotation of the 16 genes revealed several tailoring enzymes, including dioxygenases (*kpbA* and *kpbI*), PLP-dependent enzymes (*kpbB* and *kpbH*), and an FMN-dependent enzyme (*kpbJ*), as well as a hypothetical protein (*kpbL*). Interestingly, four genes, *kpbN* (glutathione *S*-transferase); *kpbP* (γ-glutamyl transferase); *kpbK* (dipeptidase); *kpbF* (ACC-synthase) were found within the *kpb* cluster, resembling gene sets found in the BGCs of gliotoxin (*gliG*, *gliK*, *gliJ*, *gliI*)^27,28^ and aspirochlorine (*aclG*, *aclK*, *aclJ*, *aclI*)^26^. In gliotoxin and aspirochlorine biosynthesis, the four enzymes generate a dithiol intermediate that is subsequently oxidized by an FAD-dependent dithiol oxidase (*gliT*, *aclT*) to form the characteristic transannular disulfide bond. In contrast, the *kpb* cluster contains no gene encoding a dithiol oxidase, and sulfur is absent from the structure of KCP. Consequently, whether these four enzymes play a role in KCP biosynthesis remains unclear, and predicting the initial step or overall pathway of KCP biosynthesis from gene functions alone proved extremely challenging.

### Comparative LC-MS/MS profiling of *E. shearii* to identify the biosynthetically related compounds of KCP

The discovery of KCP analogues in the producing fungus *E. shearii* could provide valuable insights into the biosynthetic pathway of KCP. However, even KCP itself is produced only in trace amounts from natural sources, and no intermediates or metabolites have been reported during the three decades since its isolation.^3^ These observations suggest that, even if KCP analogues exist in *E. shearii*, they are present at extremely low levels, rendering activity-guided screening impractical. We therefore employed a mass spectrometry-based approach to search for related compounds.

We first examined the MS/MS fragmentation pattern of KCP and identified a representative fragment ion at *m/z* 225.0868 (C_10_H_13_N_2_O_4_^+^, calcd. 225.0870) (Figure 3A). Although multiple structural possibilities remain, this fragment ion corresponds to a substructure lacking an aromatic ring and was considered a diagnostic marker for KCP analogues.

**Figure 3.**
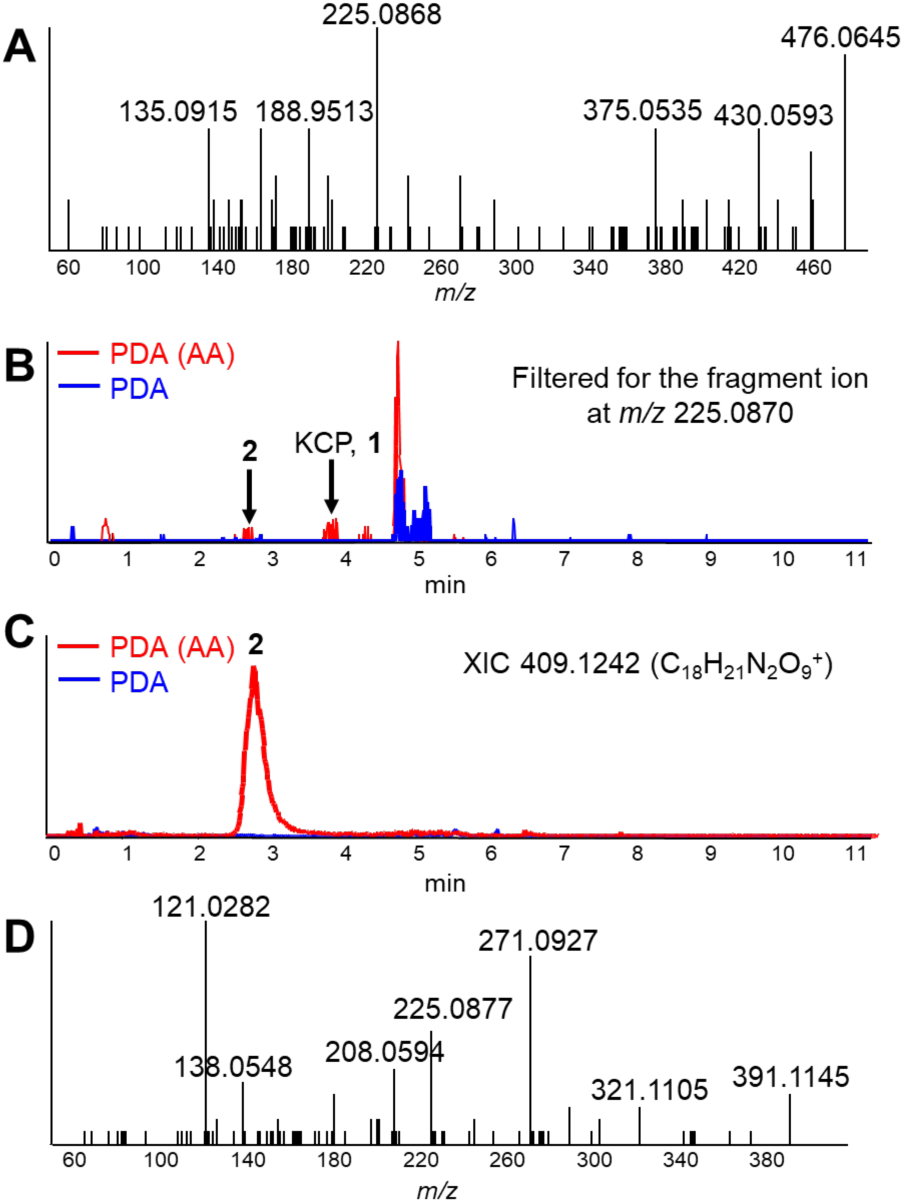
LC-MS/MS profiling for the identification of KCP analogues in *E. shearii*. (A) MS/MS fragment pattern of KCP. (B) Overlaid chromatograms filtered for the fragment ion at *m*/*z* 225.0870. Red, extract from KCP-producing condition; blue, extract from the nonproducing condition. (C) Overlaid XICs at *m/z* 409.1242 corresponding to compound **2** obtained from the two conditions. Colors as in (B). (D) MS/MS fragmentation pattern of compound **2**.

Two extracts were then subjected to LC-MS/MS analysis: one obtained from *E. shearii* cultured in PDA medium supplemented with adenine and L-arginine (KCP-producing condition), and the other from cultures without supplementation (KCP-nonproducing condition). LC-MS/MS analyses were performed on a SCIEX Q-TOF mass spectrometer operated in data-dependent acquisition (DDA) mode with a scan cycle time of 0.61 s. In each cycle, an MS^1^ full scan (*m/z* 100–1000) was first acquired, and up to 10 precursor ions were then automatically selected in order of intensity and fragmented to obtain MS/MS spectra. By overlaying the two datasets and applying fragment ion filtering at *m/z* 225.0870 (C_10_H_13_N_2_O_4_^+^) using the filtering function in SCIEX OS software (Figure 3B), we identified precursor ions containing this diagnostic fragment ion, which were observed specifically under the KCP-producing condition. This approach effectively functioned as a pseudo-“comparative precursor ion scan”.

This analysis enabled the detection of compound **2**, with the molecular formula C_18_H_20_N_2_O_9_, as a potential KCP analogue (Figure 3B, 3C). The MS/MS fragment pattern of **2** displayed not only the diagnostic fragment ion at *m/z* 225.0870 (C_10_H_13_N_2_O_4_^+^) but also a 4-hydroxybenzoyl ion at *m/z* 121.0282 (C_7_H_5_O_2_^+^, calcd. 121.0284) (Figure 3D), suggesting that **2** represents an intermediate prior to dichlorination in KCP biosynthesis (Figure S1). Furthermore, we observed that **2** was largely converted into compound **3** (C_17_H_20_N_2_O_8_) when the methanol extracts were left at room temperature overnight (Figure S2). This observation suggests that **2** contains an α-keto acid moiety that is susceptible to oxidative decarboxylation.

### Isolation of 3 and Chemical Reduction of 2 to 4 and 5

We initially aimed to isolate **2**, but it underwent oxidative decarboxylation to generate **3**. We therefore focused on the isolation and structural determination of **3**. *E. shearii* was cultivated for nine days in PDA medium supplemented with adenine and L-arginine, after which the mycelium was extracted with methanol. The methanol extract was then left at room temperature for two nights to allow complete conversion of **2** to **3**. Sequential column chromatography of the extract obtained from 7.8 L culture afforded approximately 45 μg of **3** in nearly pure form. The chemical structure of **3** was determined by MS/MS fragmentation (Figure S2) and NMR spectroscopy (¹H NMR, COSY, HSQC, and HMBC), as detailed in the Supporting Information (Figure S1, S6–S9, Tables S3–S4). These data revealed that **3** is a KCP analogue lacking two chlorine atoms, in which the D-serine moiety is replaced by a carboxymethyl group (Figure 4A). Based on this structure, we hypothesized that oxidative decarboxylation occurs at the C-4 side chain and that **2** represents a dechlorinated analogue of KCP, in which the D-serine moiety is replaced by α-keto acid, pyruvate. Accordingly, **2** and **3** were designated as dechlorokaito pyruvate and dechlorokaito acetate, respectively (Figure 4A).

**Figure 4.**
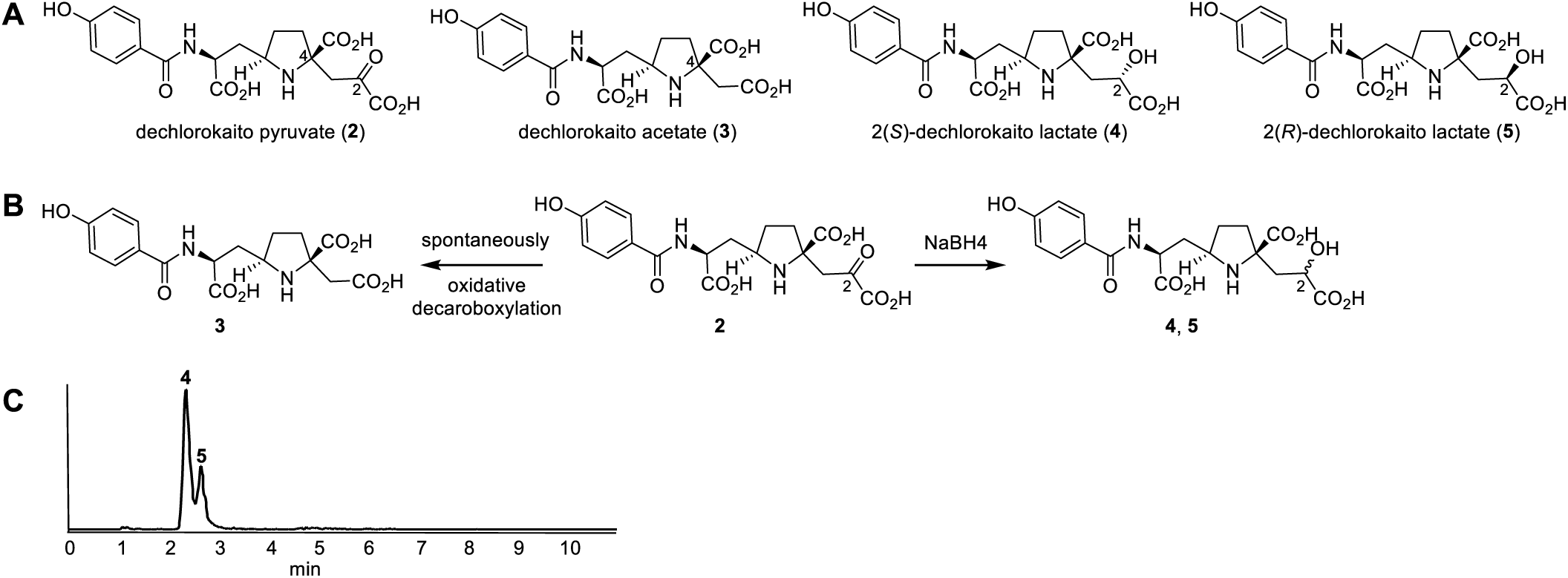
Structures of compounds **2**–**5** and their formation. (A) Structures of **2**–**5**. (B) Conversion of **2** to **3**, **4** and **5**. (C) Results of NaBH4 reduction. XIC at *m/z* 411.1398 corresponding to **4** and **5**.

To correlate the chemical structure of **2** with the predicted functions of genes in the *kpb* cluster, we focused on KpbJ, annotated as an FMN-dependent dehydrogenase. Homologous enzymes are known to act as oxidases catalyzing the conversion of α-hydroxy acids to α-keto acids.^29,30^ Thus, the α-keto acid moiety in **2** was predicted to arise from KpbJ-mediated oxidation. To examine this hypothesis, we conducted chemical reduction of **2** and used the resulting products as substrates for *in vitro* reaction. NaBH₄ reduction of **2** gave two products, compound **4** and compound **5**, in a 2:1 ratio (Figure 4B, 4C). Reduction of mycelium extracts, followed by purification through multiple chromatographic steps, ultimately yielded 87 μg of **4** and 41 μg of **5** from 26 L of culture. NMR analyses (Figures S10–S19, Tables S3–S4) confirmed that both **4** and **5** are reduced forms of **2**, in which the ketone group at C-2 is converted to a hydroxymethine, and that they are diastereomers differing at this stereocenter. Detailed spectroscopic and conformation analyses revealed that **4** possesses the *S* configuration at C-2, whereas **5** has the *R* configuration (see Supplementary Information, Figure S1). Accordingly, **4** and **5** were designated as 2(*S*)-dechlorokaito lactate and 2(*R*)-dechlorokaito lactate, respectively (Figure 4A).

### *In vitro* enzymatic reaction

Based on the chemical structures of **2**–**5**, we predicted the involvement of four enzymes, KpbJ, KpbB, KpbI, and KpbM in the biosynthetic steps leading to KCP, although the sequence of these reactions was initially unclear. We hypothesized that the FMN-dependent dehydrogenase KpbJ would catalyze oxidation of **4** or **5** to **2**; the ornithine aminotransferase KpbB would introduce an amino group at C-2; α-ketoglutarate-dependent dioxygenase KpbI would mediate hydroxylation at C-3; and the flavin-dependent halogenase KpbM would catalyze dichlorination of the 4-hydroxybenzoyl moiety. All four candidate enzymes were heterologously expressed in *E. coli* and purified using Ni-affinity chromatography (Figure S3, Tables S1–S2).

We first conducted *in vitro* assays with KpbJ using **4** or **5** as substrates. LC-MS analysis revealed that neither **2** nor its oxidatively decarboxylated product **3** was formed, indicating that KpbJ is not responsible for the oxidation of **4** or **5** to **2**. We next tested **4** or **5** as substrates with the combined addition of KpbB, KpbI, and KpbM. LC-MS analysis after overnight incubation revealed that KCP was generated in both cases (Figure 5, trace d and e), suggesting that these three enzymes together convert **4** and **5** to KCP. Moreover, the stereochemistry at C-2 did not influence KCP formation, presumably because the hydroxy group at this position is oxidized to an α-keto acid during the process.

**Figure 5.**
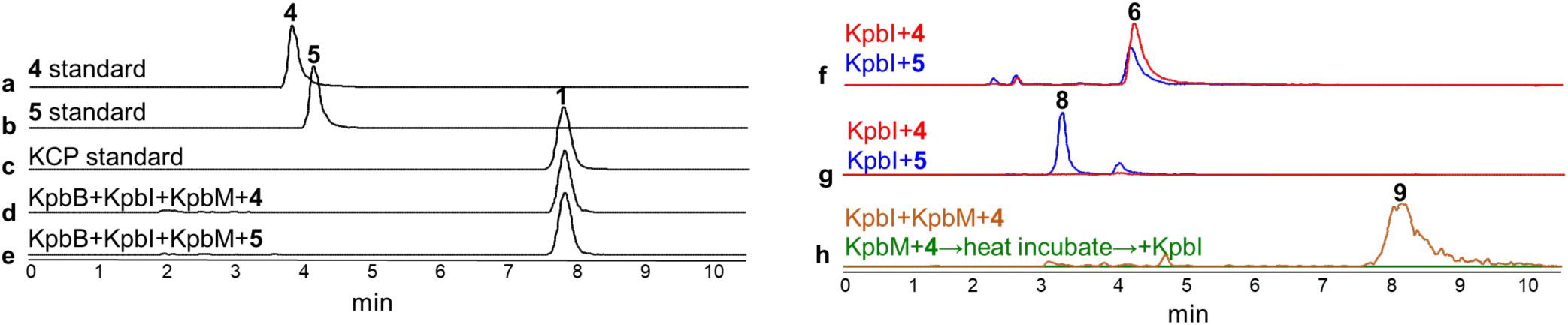
Results of *in vitro* enzyme assays. Traces **a** and **b** show XICs at *m/z* 411.1398 derived from standards **4** and **5**, respectively. Traces **c**–**e** show XICs at *m/z* 494.0728 derived from KCP (**1**) standard (**c**) and from the reaction mixtures using substrate **4** (**d**) and substrate **5** (**e**). Trace **f** shows overlaid XICs at *m/z* 397.1242 from the reaction mixtures using substrates **4** (red) and **5** (blue). Trace **g** shows overlaid XICs at *m/z* 427.1347 from the reaction mixtures using substrates **4** (red) and **5** (blue). Trace **h** shows overlaid XICs at *m/z* 465.0462 from the one-step reaction mixture (brown) and the sequential two-step reaction mixture with KpbI added afterward (green).

To examine the function of each enzyme, we carried out an *in vitro* enzymatic assay using KpbI alone with **4** or **5** as the substrate. Because the substrate concentration was 20 µM, isolation and structural determination of the products were not feasible. Instead, LC-MS analysis revealed the formation of compound **6** (C_17_H_20_N_2_O_9_) in both reactions (Figure 5, trace f). This finding indicates that KpbI catalyzes C-3 hydroxylation and subsequent oxidation of the C-2 hydroxy group to yield dechloro hydroxypyruvate (**7**), which then undergoes oxidative decarboxylation to form dechloro hydroxyacetate (**6**) (Figure 6). Further analysis revealed that the reaction with **5** also generated compound **8** (C_18_H_22_N_2_O_10_), assigned as 3-hydroxy-dechlorokaito lactate, which is consistent with hydroxylation at C-3 prior to C-2 oxidation (Figure 5, trace g). These data demonstrate that KpbI catalyzes an unprecedented two-step oxidation: hydroxylation at C-3 followed by oxidation of the C-2 hydroxyl group to a ketone. Notably, KpbI was tolerant of the stereochemistry at the C-2, as both **4** and **5** were converted.

**Figure 6.**
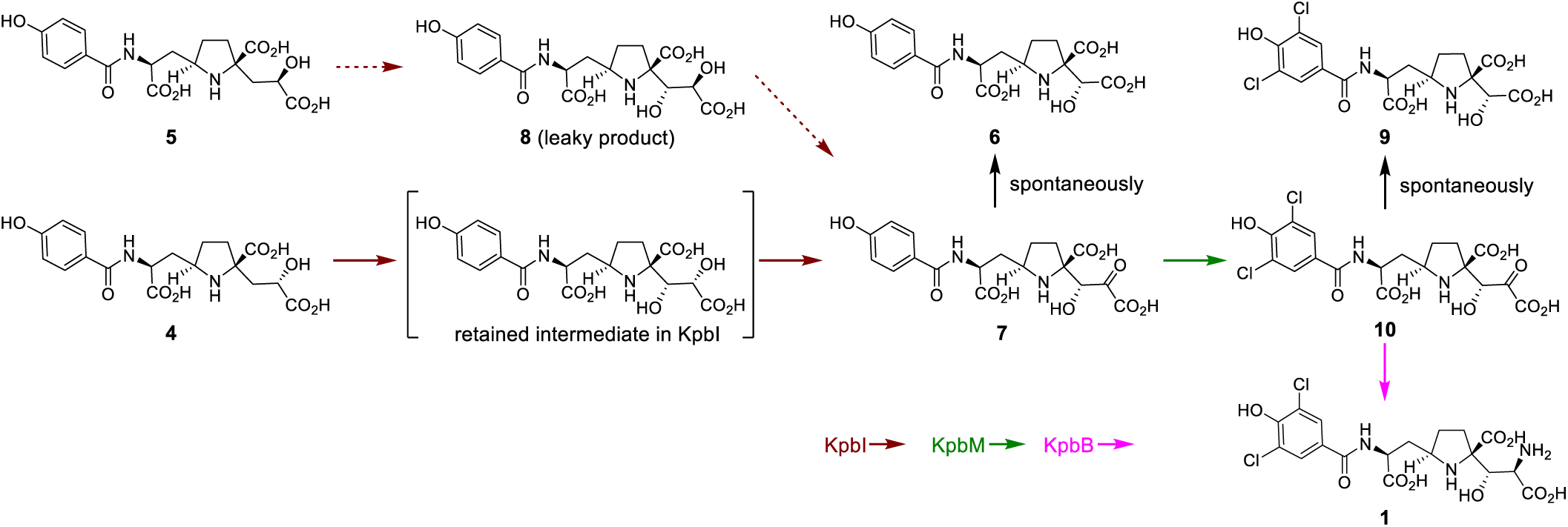
Proposed late-stage biosynthetic pathway of KCP. The dotted brown arrow from **5** to **7** represents a reaction that can occur; however, **5** is unlikely to be the genuine substrate of KpbI.

Within the α-ketoglutarate-dependent dioxygenase family, OrfP from streptolidine biosynthesis is the only other enzyme reported to catalyze hydroxylation at two distinct positions of L-arginine.^31^ OrfP hydroxylates L-arginine first at the C-3 position and subsequently at the C-4 position. In that case, a leaky intermediate (3-hydroxy-arginine) is released as a minor product after the first hydroxylation step. If compound **8** represents a leaky intermediate of KpbI, the fact that **4** is converted directly to **6** without accumulation of **8** suggests that **4** is the true substrate of KpbI. This interpretation is further supported by the higher yield of **6** when **4** was used as the substrate (Figure 5, trace f). Moreover, oxidation reactions that convert a hydroxy group into a ketone, as observed in the second step of KpbI, are exceedingly rare within the α-ketoglutarate-dependent dioxygenase family. To the best of our knowledge, thymine-7-hydroxylase is the only known enzyme reported to catalyze hydroxyl-to-carbonyl conversion, and even in that case, the reaction occurs as part of a sequential three-step process involving hydroxylation, aldehyde formation, and carboxylation.^32^ Thus, the KpbI-mediated two-step oxidation consisting of C–H hydroxylation followed by hydroxyl-to-carbonyl conversion at an adjacent position is unprecedented among this family. Interestingly, neither compound **2** nor its decarboxylated product **3** was detected in the KpbI reactions.

Although definitive evidence is not yet available, a plausible hypothesis is that the accumulation of **2** results from the KpbI-catalyzed two-step oxidation being the rate-limiting step in KCP biosynthesis. In this scenario, an endogenous enzyme with lactate dehydrogenase-like activity may act on **4** more rapidly than KpbI, converting it into **2**, which then accumulates as a shunt metabolite.

To clarify the order of subsequent steps leading KCP from **4**, we examined coupling reactions. The reaction of KpbI with KpbB using **4** as the substrate in the absence of KpbM yielded only **6** and no compound corresponding to the nonchlorinated form of KCP, suggesting that transamination occurred after dichlorination. Conversely, KpbI with KpbM in the absence of KpbB produced compound **9** (C_17_H_18_Cl_2_N_2_O_9_), designated kaito hydroxyacetate (Figure 5, trace h). To further probe the sequence, **4** was first incubated with KpbM, the enzyme was heat-inactivated, and then KpbI was added together with the metal cofactor and other required components, α-ketoglutarate and L-ascorbate. Compound **9** was not detected in this sequential assay (Figure 5, trace h), indicating that dichlorination occurred after KpbI-mediated oxidation. Consequently, these findings establish the late-stage biosynthetic pathway of KCP: **4** is first converted to **7** by dioxygenase KpbI, then to kaito hydroxypyruvate **10** by halogenase KpbM, and finally to KCP by aminotransferase KpbB (Figure 6).

### Isotope tracing experiment with [^13^C_6_]-L-arginine using *E. shearii*

Although transamination by KpbB using L-ornithine as the amino group donor is a typical reaction, it is important to examine the metabolic fate of L-ornithine following its role in this step. Upon deamination, L-ornithine generates L-glutamate-γ-semialdehyde, which is known to spontaneously cyclize to pyrroline-5-carboxylate (P5C, Figure S5). *In vivo*, P5C has been reported to undergo Mannich-type C–C bond formation with nucleophiles via its electrophilic iminium moiety.^33^ Based on this, we hypothesized that the proline moiety of KCP may be derived from P5C and that the C–C bond between C-7 and C-8 could be formed in this manner, although the identity of the nucleophilic partner remains unknown.

To test this hypothesis, *E. shearii* was cultivated in KCP-producing medium in which the L-arginine was replaced with [^13^C_6_]-L-arginine. The *kpb* cluster contains *kpbG*, annotated as an arginase converting L-arginine to L-ornithine, suggesting that L-ornithine supply is active *in vivo*. Analysis of KCP extracted from the mycelia revealed the incorporation of [^13^C5]-labeled KCP, as expected (Figure S4). These results support the idea that L-ornithine participates in KCP biosynthesis through the following route: first, L-arginine is converted to L-ornithine by KpbG; L-ornithine then donates the amino group during the late stage of KCP biosynthesis and may subsequently be incorporated as the proline moiety through P5C formation at an earlier stage. Such recycling suggests a possible, metabolic economy in the biosynthetic design of KCP (Figure S5).

Compound **4**, identified in this study as a biosynthetic intermediate, has a lactate unit attached to the α-position of the proline moiety. The formation of the C–C bond at the α-position of an amino acid is reminiscent of the involvement of a PLP-dependent enzyme such as LolT in loline biosynthesis.^34^ Within the *kpb* cluster, an uncharacterized PLP-dependent enzyme KpbH is annotated as a cysteine desulfurase, and it may contribute to this C–C bond formation. In contrast, the identity of the coupling partner at the δ-position of the proline, which may be the apparent alanine moiety or another molecule, remains unsolved. Clarifying the enzymatic machinery responsible for these unique bond formations will be an important direction for future investigation.

## Conclusion

Nearly three decades after its discovery, KCP has remained an enigmatic natural product owing to its scarcity and the lack of biosynthetic insights. Unlike canonical metabolites derived from polyketides, nonribosomal peptides, or terpenoids, KCP does not belong to any known biosynthetic class. It makes genome-mining guided by scaffold-forming enzymes ineffective for the identification of BGC. By integrating genomic, transcriptomic, metabolomic, and enzymatic approaches, we have successfully identified the *kpb* cluster and elucidated critical late-stage steps in its assembly. We demonstrate that the D-serine moiety is not directly introduced as an amino acid but is constructed stepwise through an unprecedented two-step oxidation by the α-ketoglutarate-dependent dioxygenase KpbI.

Building on the mechanistic insight, isotope tracing experiments support that the proline moiety is derived from L-ornithine via P5C, suggesting a recycling strategy in which an amino donor is subsequently incorporated into the product scaffold. This strategy appears to be in line with the observed increase in KCP production upon arginine supplementation in the culture medium, lending further support to its physiological relevance. Collectively, there is little doubt that KCP biosynthesis still harbors intriguing mechanistic features. This study establishes a solid foundation for complete pathway elucidation.

## Supporting Information

The authors have cited additional references within supporting Information. [35–40]

## Notes

The authors declare no competing financial interest.

## Acknowledgements

We are grateful to Dr. Hiroyuki Watanabe (The University of Tokyo) and Dr. Kei Kudo (National Institute of Advanced Industrial Science and Technology) for their kind assistance with the NMR measurements. This work is supported by the grants from JSPS KAKENHI (22H05120 to T.K., 23H04547 to J.M., and 25K18178 to Y.M.). Y.M. was supported by a Grant-in-Aid for JSPS Postdoctoral Fellow (22KJ0725) and the Challenging Research Grant for Young Female Scientists from the Japan Society for Bioscience, Biotechnology, and Agrochemistry (JSBBA).

## General Experimental Procedures

The biochemicals and enzymes for genetic manipulation were purchased from TaKaRa Bio (Ohtsu, Japan) and TOYOBO (Osaka, Japan). Oligonucleotides used for genetic manipulation (Table S1) were purchased from Fasmac Co., Ltd. (Kanagawa, Japan). All other reagents were purchased from FUJIFILM Wako Pure Chemical Industries (Osaka, Japan), Kanto Chemicals (Tokyo, Japan), Tokyo Chemical Industry (Tokyo, Japan) and Nacalai Tesque (Kyoto, Japan) unless otherwise noted. Cells of *E. coli* were disrupted using a Branson Sonifier 250 (Emerson Japan, Tokyo, Japan). DNA manipulation was performed according to the manufacturer’s instructions. NMR spectra were acquired with a Varian NMR 600 NB CL spectrometer (Varian, CA, U.S.A.) in 0.2 mL (Symmetrical Micro Sample Tube, cat. BMS-005V, Shigemi, Hachioji, Japan) of D_2_O (deuteration degree: 99.95%). The data were referenced to residual solvent signals with resonances at *δ*_H/C_ = 3.30/49.0 ppm (remaining metanol). UHPLC–MS analysis was performed with a Nexera X3 (Shimadzu, Kyoto, Japan) coupled to an X500R QTOF system (SCIEX, Framingham, MA, USA), which was equipped with an electrospray source operating in the positive-ionization mode.

### Strains

*Escherichia coli* DH5α was used for cloning following standard recombinant DNA techniques. *E. coli* BL21(DE3) was used for expression of protein. *Eupenicillium shearii* PF1191^1^ was used for screening and isolation of KCP-related compounds, genome sequencing and RNA sequencing.

### Kaitocephalin-producing conditions

To ensure stable kaitocephalin production, we tested and optimized various culture conditions. We established the following stable kaitocephalin production conditions. Mycelia of *E. shearii* were inoculated into a liquid MPY medium of 100 mL at 30°C with shaking at 150 rpm for 18 hours. After filtration, the mycelium was inoculated into 130 mL of potato dextrose agar (PDA) medium supplemented with 0.1% of L-arginine and 0.05% of adenine. The plates were incubated at 25°C for 5–14 days. The non-producing conditions refer either to cultivation on PDA medium lacking thymine or alanine, or to cases where, despite supplementation, the mycelia were harvested before 5 days of culture.”

### Genomic DNA preparation

Genomic DNA was extracted according to the following method; the mycelia of *E. shearii* PF1191 was collected by filtration, washed with water, and dried using paper towel. The dried mycelia were frozen in liquid nitrogen and ground into a fine powder using a mortar and pestle. To the frozen powder was added extraction buffer (400 mM of Tris-HCl (pH 8.0), 500 mM of NaCl, 20 mM of ethylenediaminetetraacetic acid (EDTA) and 1% of sodium dodecyl sulfate) and the suspension was kept at room temperature for 5 min. To the suspension was added phenol: chloroform solution and the mixture was vortexed for 2 sec. After incubation at 65 °C for 60 min, the reaction mixture was centrifuged at 12000 rpm for 5 min. The supernatant was then treated with RNase at 37 °C for 90 min. To the reaction mixture was then added phenol-chloroform-isoamyl alcohol (PCI) solution. After being vortexed for 2 sec, the mixture was centrifuged at 12000 rpm for 5 min. The supernatant was transferred to a new centrifuge tube and re-extracted twice with PCI solution followed by chloroform. To the final supernatant was added cold-isopropanol and CH_3_COONa solution and genomic DNA was recovered by centrifugation at 12000 rpm for 10 min. The pellet was then washed with 70% ethanol solution and dried for 15 min. Finally, the isolated DNA was resuspended in TE buffer (10 mM of Tris-HCl (pH 8.0) and 1mM of EDTA) and stored at -20 °C for further use.

### Genome analysis

Genome sequencing and *de novo* assembly of *E. shearii* PF1191 were performed by Genome-Lead Corp. (Kagawa, Japan) using an Illumina HiSeq system (Illumina, San Diego, CA, USA) and the assembler Shovill (v1.1.1). The resulting draft genome comprised 33 contigs (≥1 kb) with a total length of approximately 34.5 Mbp. Gene prediction was then performed with Augustus,^2^ fungiSMASH,^3^ and 2ndfind.^4^ The prediction models of AUGUSTUS and 2ndfind were trained with *Aspergillus oryzae* as the reference.

### Comparative transcriptome analysis

Mycelia of Aspergillus were obtained from potato dextrose agar plates cultured under four conditions: with supplementation of adenine and L-arginine or without supplementation, and harvested either on day 2 or day 5. Total RNA was extracted using NucleoSpin RNA (Macherey-Nagel, Düren, Germany). RNA sequencing was conducted by Novogene Co. Ltd. (Beijing, China) using Illumina NovaSeq 6000 Sequencing System. The data processing was conducted as follows. Quality control of the raw RNA-seq reads was performed using FastQC (v0.11.6), and adapter removal and quality trimming was conducted with fastp (v0.23.2). The cleaned reads were then aligned to the *E. shearii* reference genome using HISAT2 (v2.2.1), generating SAM files. These files were converted to BAM format using SAMtools (v1.22). Read counts and TPM (Transcripts Per Million) values were calculated using StringTie (v2.2.2) from BAM files derived from four different conditions, with a GTF annotation file used as a guide. Z-scores were then calculated for each gene across the conditions based on TPM values, and a heatmap was generated in R using these z-scores to visualize expression patterns.

### Data availability

The nucleotide sequences of *kpbA*–*kpbP* sequence has been deposited in the DNA Data Bank of Japan (DDBJ) with the accession number LC896246.

### Comparative LC-MS/MS profiling for the identification of the identification of KCP analogues

Mycelia were collected from 130 mL of potato dextrose agar media cultured for 9 days under both KCP-producing conditions (with adenine and arginine supplementation) and non-producing conditions (without supplementation). The collected mycelia were extracted with 3 mL of methanol. The extracts were centrifuged at 12,000 × g for 5 minutes, and the supernatants were loaded onto Cosmosil 140C_18_-OPN column (0.5 mL, Nacalai Tesque, Kyoto, Japan) and eluted with methanol. An aliquot of each eluate was subjected to HR-LC-MS/MS. The analytical conditions were LC: column, CAPCELLPAK C18 IF column (2.0 × 50 mm, 2 µm, Osaka Soda, Osaka, Japan); flow rate, 0.3 mL/min; solvent system: solvent A (water including 0.1% formic acid) and solvent B (acetonitrile including 0.1% formic acid); gradient program: 0–3 min 5% B, 3–8 min 5–90% B, 8–10 min 90% B, 10–11 min 5% B, 11–14 min 5% B; column temperature: 40°C. The MS detection was performed using an X500R QTOF system. The parameters for ESI(+)-DDA analysis were as follows: MS1 mass range, *m/z* 100–1000; MS2 mass range, *m/z* 50– 1000; MS1 accumulation time, 250 ms; MS2 accumulation time, 30 ms; maximum candidate ions, 10; ion source temperature, 350°C; and CAD gas, 7; ion source gas 1, 60 psi; ion source gas 2, 60 psi; curtain gas, 30 psi; spray voltage, 5500 V; declustering potential, 50 V; and collision energy, 35 eV. Product ion filtering was conducted using the Explorer function in SCIEX OS. Formula Finder (SCIEX OS) was used to determine the molecular formulae of the detected ions.

### Purification of dechlorokaito acetate (3)

*E. shearii* was cultured on one plate containing 130 mL of potato dextrose agar medium supplemented with 0.1% of L-arginine and 0.05% of adenine at 25°C for 9 days. The mycelium collected from 10–20 plates were homogenized with 200 mL of methanol twice. The extracts were centrifuged at 12,000 × g for 5 minutes, and the supernatant containing **2** was filtered using filter paper and left at room temperature for two days to enable the spontaneous oxidative decarboxylation from **2** to **3**. The filtrate was evaporated and redissolved in 20 mL of water. The extract was loaded onto an activated charcoal column (30 mL). The column was washed with water (200 mL) and eluted with aqueous acetone (200 mL). The eluate was evaporated under vacuum and dissolved in H_2_O-HCOOH (100:0.1, *v*/*v*, 30 mL). The solvent was loaded onto a Cosmosil 140C_18_-OPN column (6 mL) pre-equilibrated with H_2_O-HCOOH (100:0.1, *v*/*v*). After washing the column with the same solvent (20 mL), the solvent H_2_O-MeOH-HCOOH (90:10:0.1, *v*/*v*/*v*, 30 mL) was supplied to the resin. The eluate was concentrated and applied to an InertSustain C18 column (7.6 × 250 mm, 5 µm, GL Sciences, Tokyo, Japan) with H_2_O-MeOH-HCOOH (86:14:0.1, *v*/*v*/*v*). Fractions containing **3** were then concentrated and applied to a Mightysil RP-18GP column (4.6 × 250 mm, 5 µm, Kanto Chemical, Tokyo, Japan) with H_2_O-MeOH-HCOOH (90:10:0.1, *v*/*v*/*v*). Subsequently, fractions containing **3** were concentrated again and applied to a Mightysil RP-18GP column (4.6 × 250 mm, 5 µm) with H_2_O-MeOH-HCOOH (93:7:0.1, *v*/*v*/*v*). Compound **3** (32 µg) was obtained in almost pure form 60 plates, corresponding to 7.8 L of the agar medium. Since the amount of the isolated compound was insufficient for accurate weighing and no authentic standard was available, its quantity was estimated by LC-MS using L-tyrosine as an external standard. [M+H]^+^ **2**: *m/z* 409.1240, calcd. for C_18_H_21_N_2_O_9_^+^ 409.1242. [M+H]^+^ **3**: *m/z* 381.1294, calcd. for C_17_H_21_N_2_O_8_^+^ 381.1292.

### Reduction of 2 affording 4 and 5, and their purification

The procedures from cultivation to filtration using filter paper were the same as those for the purification of **3**. To the methanol filtrate (200 mL), NaBH_4_ (100 mg, 2.6 mmol) was added, and stirred for 30 min at room temperature, affording **4** and **5** from **2**. The reaction mixture was evaporated redissolved in 20 mL of water. The procedures of purification using activate charcoal column, cosmosil140C_18_-OPN, and InertSustainC18 were the same as those for the purification of **3**. Compounds **4** and **5** were eluted separately. Next, the eluates containing **4** and **5** were each concentrated and applied to a Mightysil RP-18 GP column (4.6 × 250 mm, 5 µm) with H_2_O-MeOH-HCOOH (93:7:0.1, *v*/*v*/*v*). The eluate containing **4** was concentrated and applied to a Mightysil RP-18 GP Aqua column (4.6 250 mm, 5 µm) with H_2_O-MeCN-HCOOH (96:4:0.1, *v*/*v*/*v*). As a result, almost pure **4** (87 µg) was obtained. Next, the eluate containing **5** from purification with InertSustain C18 column as above was concentrated and applied to a Mightysil RP-18 GP column (4.6 250 mm) with H_2_O-MeOH-HCOOH (93:7:0.1, *v*/*v*/*v*). The eluate was concentrated and applied to a Mightysil RP-18 GP Aqua column (4.6 250 mm, 5 µm) with H_2_O-MeCN-HCOOH (96:4:0.1, *v*/*v*/*v*). The eluate containing **5** was concentrated and further purified by cation exchange-reverse phase (CE-RP) chromatography using a TCI Dual ODS CX15 column (4.6 × 250 mm, 5 µm) with H_2_O-MeCN-HCOOH (85:15:0.1, *v*/*v*/*v*). Finally, **4** (41 µg) was obtained in almost pure form. [M+H]^+^ **4**: *m/z* 411.1401, **3**: *m/z* 411.1402, calcd. for C_18_H_23_N_2_O_9_^+^ 411.1398.

### Conformation analysis of 4 and 5

Conformational searches were initially performed using Avogadro (version 1.2.0) to generate possible conformers of **4** and **5**. The lowest-energy conformers were subsequently subjected to geometry optimization using Chem3D (PerkinElmer, version 23.1.1) with the MM2 force field, in order to predict the most stable conformations. The interatomic distances were predicted using the measurement tools implemented in Chem3D.

### Cloning

Plasmid constructs containing codon-optimized synthetic genes for *kpbB*, *kpbI*, and *kpbJ* for *E. coli*, cloned into pET-28a(+) with an N-terminal His_6_tag, were purchased from Twist Bioscience (San Francisco, CA, USA). A synthetic gene fragment encoding *kpbM* was also purchased from Twist Bioscience. *KpbM* gene was amplified by PCR using KOD One PCR Master Mix (TOYOBO, Osaka, Japan) to incorporate the appropriate HiFi DNA assembly overhangs. The *fre* gene, encoding an NAD(P)H-flavin reductase, was also amplified by PCR using an *E. coli* colony as template to incorporate the appropriate HiFi DNA assembly overhangs. Each PCR product was individually cloned into the pHis8 vector,^5^ which had been digested with BamHI and HindIII using NEBuilder HiFi DNA Assembly Master Mix (New England Biolabs, Ipswich, MA, USA), yielding the plasmids pHis8-*kpbM* and pHis8-*fre*, respectively.

### Purification of recombinant proteins from *E. coli* BL21(DE3)

After introducing each plasmid into *E. coli* BL21(DE3), each transformant was cultivated in 200 mL TB medium (tryptone 1.2%, yeast extract 2.4%, glycerol 0.56%, and K_2_HPO_4_ 0.23% supplemented with kanamycin) at 37°C for 5h until the cells reached an OD_600_ of ∼0.7. After cooling the culture on ice for 10 min, β-D-thiogalactopyranoside (IPTG) was added to the culture at a final concentration of 200 µM to induce gene expression, and the culture was further incubated at 18 °C for 12 h. The cells were harvested by centrifugation. Five recombinant proteins, KpbB, KpbI, KpbJ, KpbM, and Fre were purified using Ni-affinity chromatography under the same general protocol. KpbI, KpbM, and Fre were purified using standard wash buffer (50 mM NaH_2_PO_4_ (pH 8.0), 300 mM NaCl, 20 mM imidazole-HCl (pH 8.0), and 10% glycerol) and elution buffer (50 mM NaH_2_PO_4_ (pH 8.0), 300 mM NaCl, 250 mM imidazole-HCl (pH 8.0), and 10% glycerol). For KpbB, the same purification procedure was followed, but both buffers were supplemented with 0.5 mM pyridoxal 5′-phosphate (PLP). Similarly, the buffers used for KpbJ were supplemented with 0.5 mM flavin mononucleotide (FMN). After centrifugation, the cells were resuspended in 20 mL of wash buffer and lysed by sonication on ice. Cell debris was removed by centrifugation at 4 °C (30,000 × g, 20 min). The supernatant was loaded onto 2 mL of Ni-NTA Superflow Resin (Qiagen, Tokyo, Japan), and the resin was washed with 20 mL of wash buffer. The bound proteins were eluted with 20 mL of elution buffer. Each purified protein sample was analyzed by SDS‒PAGE. Finally, the proteins were concentrated using a Vivaspin 20 ultrafiltration unit (Sartorius, Göttingen, Germany), and storage buffer (20 mM Tris-HCl (pH 8.0), 100 mM NaCl, 1 mM DTT, and 20% glycerol) was added. The resulting protein solutions were stored at −80°C.

### In vitro assay

<REACTION of KpbJ>

The reaction was performed in a total volume of 100 μL of buffer comprising 100 mM potassium phosphate buffer (pH 7.5), 20 μM FMN, 20 mg/L catalase, 20 μM **4** or **5**, and 2 μM KpbJ. The reaction mixture was incubated at 25 °C overnight. After incubation, the reaction was quenched by adding 100 μL of methanol. After centrifugation, the supernatant was subjected to HR-LC-MS analysis using a X500R mass spectrometer. LC was performed using a column, Mightysil RP-18GP Aqua (2.0 × 150 mm, 5 µm, Kanto Chemical) with H_2_O-MeCN-HCOOH (93:7:0.1, *v*/*v*/*v*). The flow rate was 0.2 mL/min. The analytical conditions were used in all other *in vitro* assay.

<Reaction of KpbB, KpbI, and KpbM>

The reaction was performed in a total volume of 100 μL of buffer comprising 100 mM potassium phosphate buffer (pH 7.5), 25 mM ascorbic acid, 50 μM Fe(NH_4_)_2_(SO_4_)_2_, 1 mM α-ketoglutarate, 1 mM L-ornithine, 10 mM PLP, 1 μM FAD, 2.5 mM NADH, 1 mM MgCl_2_, 20 μM **4** or **5**, and 2 μM KpbB, KpbI, KpbM. The reaction mixture was incubated at 25 °C overnight. After incubation, the reaction was quenched by adding 100 μL of methanol. After centrifugation, the supernatant was subjected to HR-LC-MS analysis.

<REACTION of KpbI>

The reaction was performed in a total volume of 100 μL of buffer comprising 100 mM potassium phosphate buffer (pH 7.5), 25 mM ascorbic acid, 50 μM Fe(NH4)2(SO4)2, 1 mM α-ketoglutarate, 20 μM **4** or **5** and 2 μM KpbI. The reaction mixture was incubated at 25 °C overnight. After incubation, the reaction was quenched by adding 100 μL of methanol. After centrifugation, the supernatant was subjected to HR-LC-MS analysis.

<REACTION of KpbB and KpbI>

The reaction was performed in a total volume of 100 μL of buffer comprising 100 mM potassium phosphate buffer (pH 7.5), 25 mM ascorbic acid, 50 μM Fe(NH4)2(SO4)2, 1 mM α-ketoglutarate, 1 mM L-ornithine, 10 mM PLP, 20 μM **4**, and 2 μM KpbB, KpbI. The reaction mixture was incubated at 25 °C overnight. After incubation, the reaction was quenched by adding 100 μL of methanol. After centrifugation, the supernatant was subjected to HR-LC-MS analysis.

<REACTION of KpbI and KpbM>

One step reaction: The reaction was performed in a total volume of 100 μL of buffer comprising 100 mM potassium phosphate buffer (pH 7.5), 25 mM ascorbic acid, 50 μM Fe(NH4)2(SO4)2, 1 mM α-ketoglutarate, 1 μM FAD, 2.5 mM NADH, 1 mM MgCl2, 20 μM **4**, and 2 μM KpbI, KpbM. The reaction mixture was incubated at 25 °C overnight. After incubation, the reaction was quenched by adding 100 μL of methanol. After centrifugation, the supernatant was subjected to HR-LC-MS analysis.

Two step reaction: The first reaction of KpbM was performed in a total volume of 100 μL of buffer comprising 100 mM potassium phosphate buffer (pH 7.5), 1 μM FAD, 2.5 mM NADH, 1 mM MgCl2, 20 μM **4**, and 2 μM KpbM. The reaction mixture was incubated at 25 °C overnight. After incubation, the reaction was quenched by heating at 95 °C for 5 min and centrifuged. The supernatant was collected and applied to the second reaction. The reaction was performed by adding 25 mM ascorbic acid, 50 μM Fe(NH4)2(SO4)2, and 1 mM α-ketoglutarate. The reaction mixture was incubated at 25 °C overnight. After incubation, the reaction was quenched by adding 100 μL of methanol. After centrifugation, the supernatant was subjected to HR-LC-MS analysis.

### Labeling experiment with [^13^C6]-L-arginine using *E. shearii*

Mycelia of *E. shearii* were inoculated into a liquid MPY medium of 100 mL at 30°C with shaking at 150 rpm for 18 hours. After filtration, the mycelium was inoculated into 20 mL of potato dextrose agar medium supplemented with 0.1% of [^13^C_6_]-L-arginine and 0.05% of adenine. The plates were incubated at 25°C for 9 days. The collected mycelia were extracted with 2 mL of methanol. The extracts were centrifuged at 12,000 × g for 5 minutes, and the supernatants were loaded onto Cosmosil 140C_18_-OPN column (0.5 mL) and eluted with methanol. An aliquot of each eluate was subjected to HR-LC-MS/MS. The analytical conditions were the same as those of “Comparative LC-MS/MS profiling for the identification of biosynthetically related compounds of KCP” section.

### Structural analysis of 3, 4, and 5

The chemical structure of **3** was elucidated by MS/MS analysis and NMR experiments, including ¹H NMR, COSY, HSQC, and HMBC. First, the presence of a 4-hydroxybenzoyl moiety was inferred from an MS/MS fragment ion at *m/z* 121.0281, corresponding to C_7_H_5_O_2_^+^ (calcd. 121.0284) (Figure S2), and from two aromatic doublet signals (H-12/H-16 and H-13/H-15) observed in the ¹H NMR spectrum. Next, the COSY correlations of H-5/H-6, H-6/H-7, and H-8/H-9 and HMBC correlations of H-5/C-1, H-6/C-8, H-7/C-8, and H-9/C-17 suggested that **3** contains a proline ring disubstituted at both α-and δ-carbons like KCP. The side chain at C-7 (δ-carbon) would be identical to that of KCP based on the similarity of the carbon and proton chemical shifts at positions 8, 9, and 17 and 2D NMR correlations. On the other hand, the side chain at C-4 (α-carbon) appeared to differ from that of KCP. In compound **3**, nonequivalent methylene protons are present at C-3, and the chemical shift of this carbon (40.0 ppm) is clearly distinct from that of the hydroxymethine carbon at C-3 in KCP (71.4 ppm). Furthermore, as shown in the HMBC spectrum, correlations were observed from H-3 to C-2 (174.2 ppm), indicating that the carboxyl group is attached to the 3-CH₂ position. In this way, the planar structure of **3** was determined, and all proton and carbon signals were assigned as summarized in Table S3. The stereochemistry of **3** could not be directly determined, as no correlation was observed in the 1D NOESY spectra due to the limited sample amount. However, comparison of the ^1^H NMR data of **3** with those of a series of chemically synthesized KCP analogues,^6,7^ including both analogues retaining the same 7*R*,9*S* configuration as KCP and those with altered stereochemistry at C-7 and C-9 indicated that **3** possesses the same stereochemistry as KCP. Specifically, in KCP itself and analogues retaining the same 7*R*,9*S* configuration, one of the nonequivalent protons of the 8-CH₂ group, located between C-7 and C-9, appeared at approximately 2.5 ppm and was clearly separated from signals of the proline ring (H-5, H-6). This distinctive C-8 methylene proton signal was also observed for **3**, supporting its assignment as having the same relative stereochemistry as KCP. In contrast, this feature was absent in the C-7 or C-9 stereoisomers. Since the absolute configuration of KCP has been established, **3** was consequently assigned the same absolute configuration (7*R*,9*S*). Although no spectral data were available to determine the stereochemistry at C-4 in this stage, it was inferred from the analysis of compound **4**, which is described below.

NMR experiments including ^1^H, COSY, HSQC, and HMBC confirmed that both **4** and **5** are reduced forms of **2**, in which the ketone group at C-2 is converted to hydroxymethine (*δ*_H_ 4.30, *δ*_C_ 68.0 for **4**; *δ*_H_ 4.09, *δ*_C_ 69.0 for **5**), and that they are diastereomers differing in the stereochemistry at this position. In the minor isomer **5**, no signals were detected in the 1D NOESY spectrum because of the limited amount of sample. In contrast, in the major isomer **4**, key NOE correlations were observed, which enabled assignment of its stereochemistry. Specifically, H-5α which showed NOE correlation with H-7, also correlated with H-2, supporting that the lactic acid side chain at C-4 is connected via the α-position (Figure S1B). Additionally, conformational analysis of **4** and **5** revealed that the H-2–H-5α distances are both 2.3 Å in the 2*S* and 2*R* configurations, whereas the H-7–H-5α distances differ markedly, being 2.6 Å in the 2*S* configuration and 3.8 Å in the 2*R* configuration (Figure S1C). In principle, the NOESY signal intensity is inversely proportional to the sixth power of the internuclear distance, and a distance of 3.8 Å is known to give only a weak signal.^8,9^ Therefore, the observation that selective irradiation of H-5α in **4** produced H-7 and H-2 signals of comparable intensity strongly suggests that **4** possesses the 2*S* rather than the 2*R* configuration. Since chemical reduction with NaBH_4_ is unlikely to alter the stereochemistry at C-4, C-7 and C-9, the stereochemistry is assumed to be identical to that in **3**. Accordingly, we accomplished the structural elucidation of **2**–**5**, including their stereochemistry (Figure S1A). NMR data of **3**–**5** were summarized in Table S3 and S4.

---

**Figure S1.**
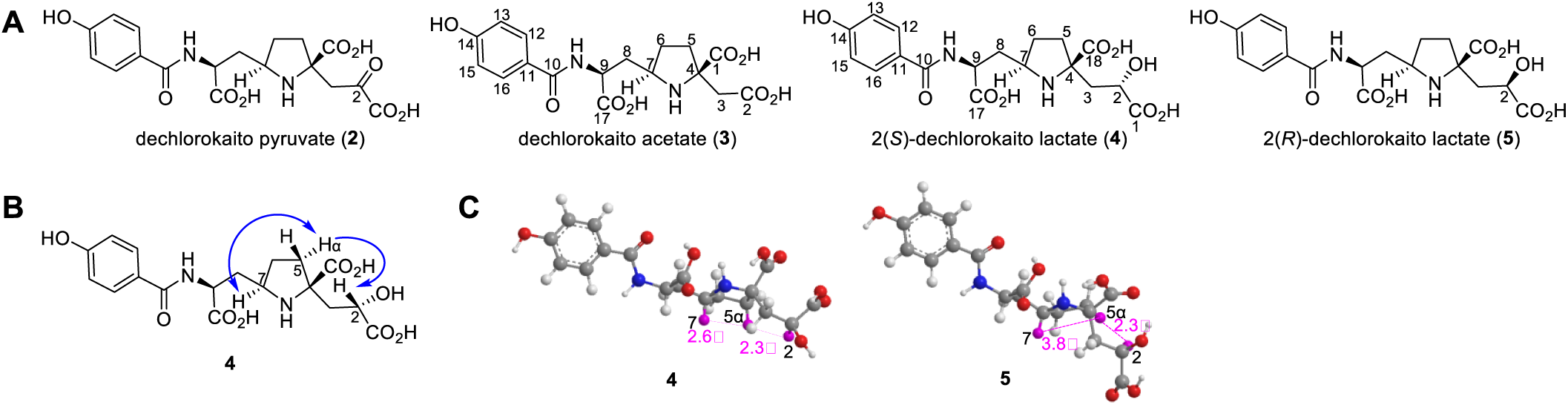
Chemical structures of **2**–**5**. (A) Chemical structures of **2**–**5**. (B) Key NOEs of **4**. (C) Calculated stable conformations of **4** and **5** and the internuclear distances between H-2–H-5α, and between H-5α–H-7. Atoms of H-2, H-5α, and H-7 are highlighted in pink.

**Figure S2.**
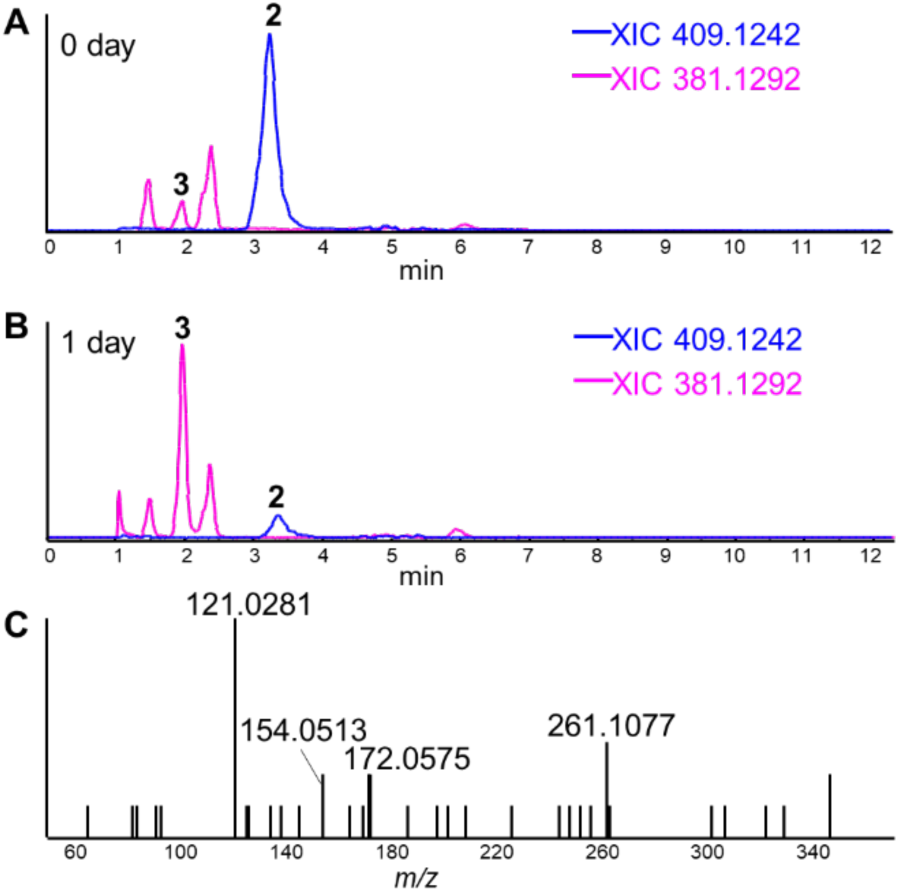
Spontaneous conversion of **2** to **3**. (A) XICs at *m/z* 409.1242 (**2**, blue) and at *m/z* 381.1292 (**3**, pink) derived from the extract sample prepared immediately after harvest. (B) XICs at *m/z* 409.1242 (**2**, blue) and at *m/z* 381.1292 (**3**, pink) derived from the extract sample left for 1 day. (C) MS/MS fragment pattern of **3**.

**Figure S3.**
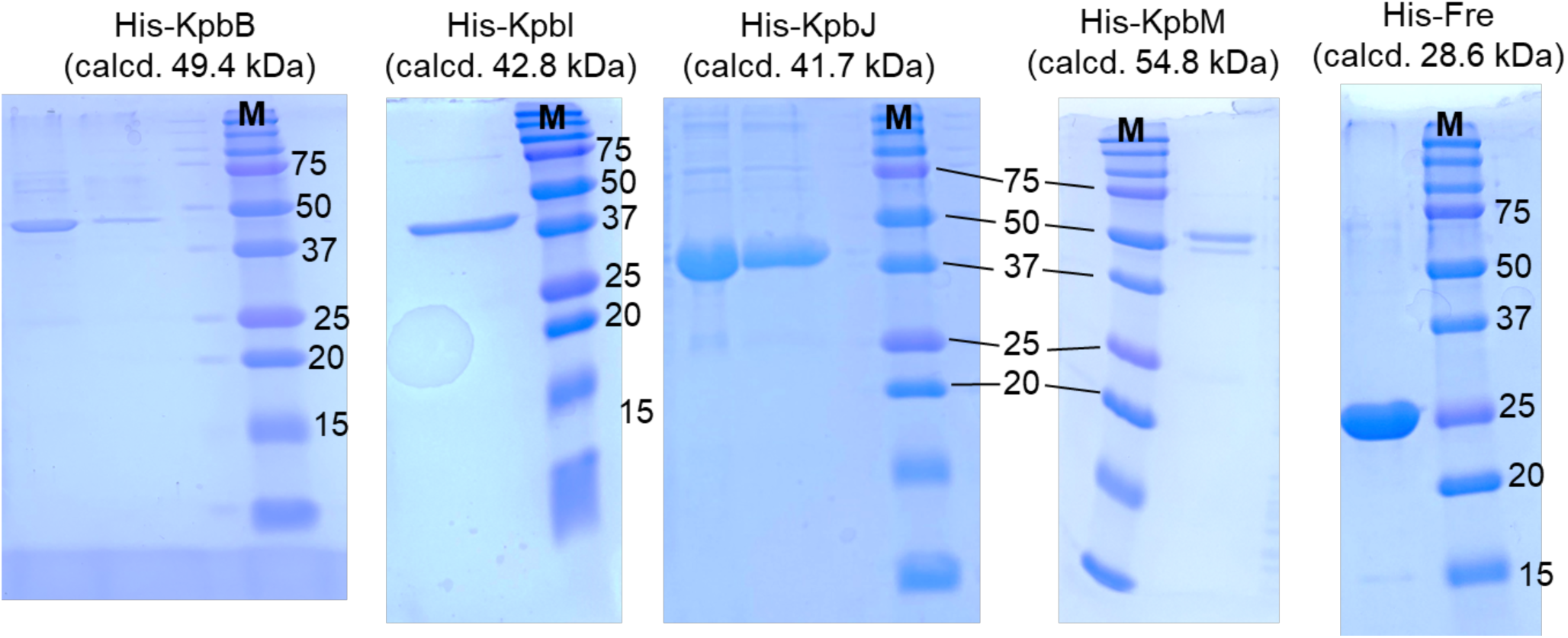
SDS‒PAGE analysis of purified recombinant proteins. SDS‒PAGE (12%) analysis of recombinant proteins. M, molecular weight marker. Proteins were produced as Histagged proteins and purified by Ni-NTA affinity chromatography using Ni-NTA Superflow Resin. The two lanes for His-KpbB and His-KpbJ represent fractions collected from the 20 mL eluate, corresponding to the first 10 mL (left) and the remaining 10 mL (right).

**Figure S4.**
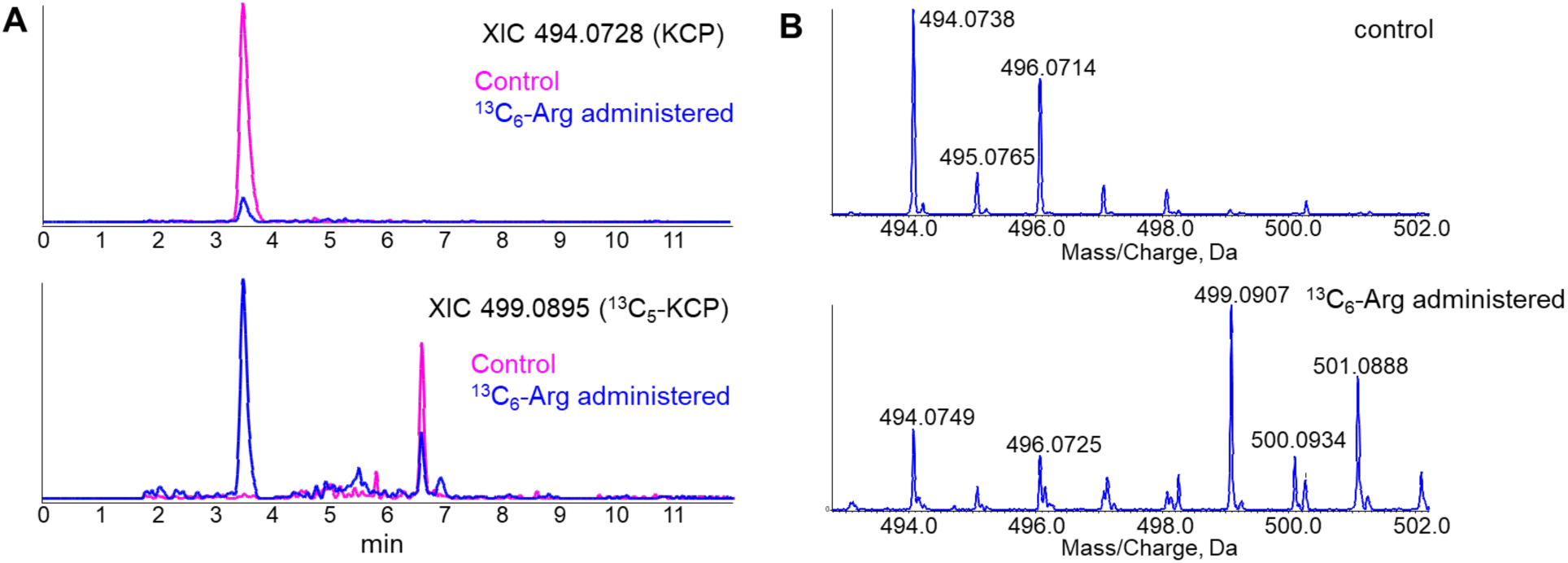
Feeding experiment with [^13^C_6_]-L-arginine. (A) XICs at *m/z* 494.0728 corresponding to KCP (top) and at *m/z* 499.0895 corresponding to [^13^C_5_]-KCP (bottom). Pink chromatogram: extracts from *E. shearii* grown on medium with adenine and L-arginine (control). Blue chromatogram: extracts from *E. shearii* grown on medium with adenine and [^13^C_6_]-L-arginine replacing L-arginine (^13^C_6_-Arg administered). The peak at 6.6 min in the bottom chromatogram, based on HRMS and the predicted molecular formula, represents an impurity unrelated to KCP. (B) The isotope patterns of KCP from control extracts (top) and from ^13^C_6_-Arg administered extracts (bottom).

**Figure S5.**
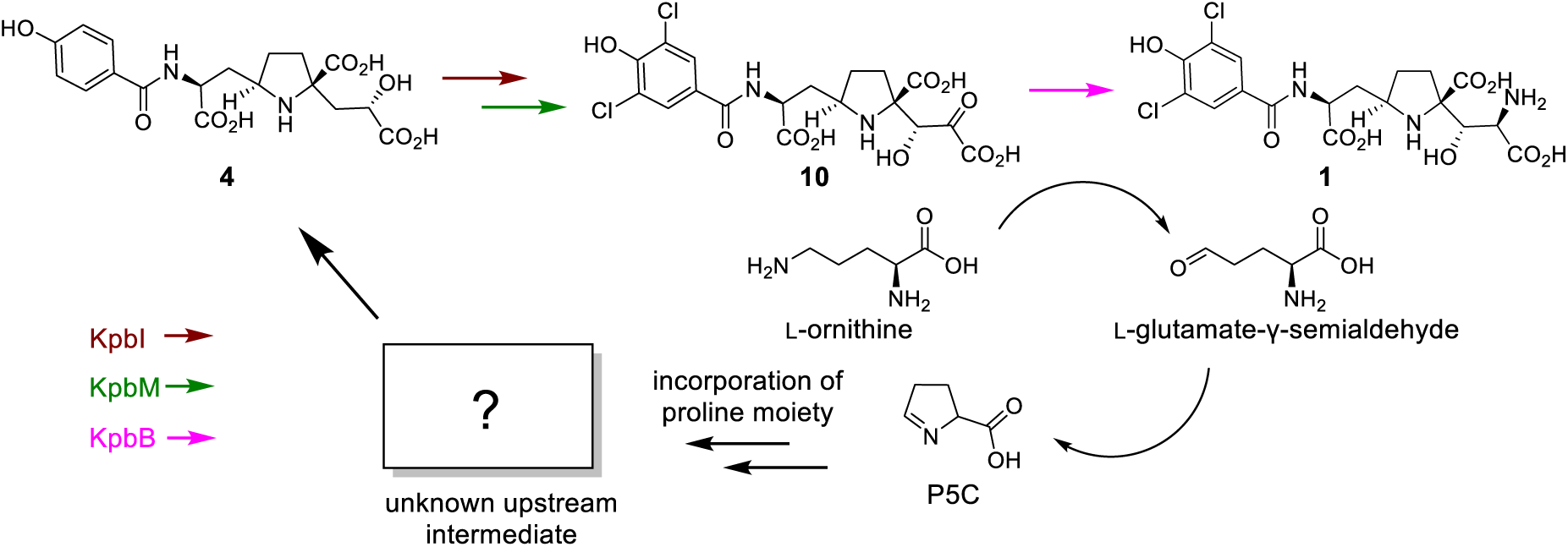
Predicted biosynthetic pathway that efficiently utilizes L-ornithine.

**Figure S6.**
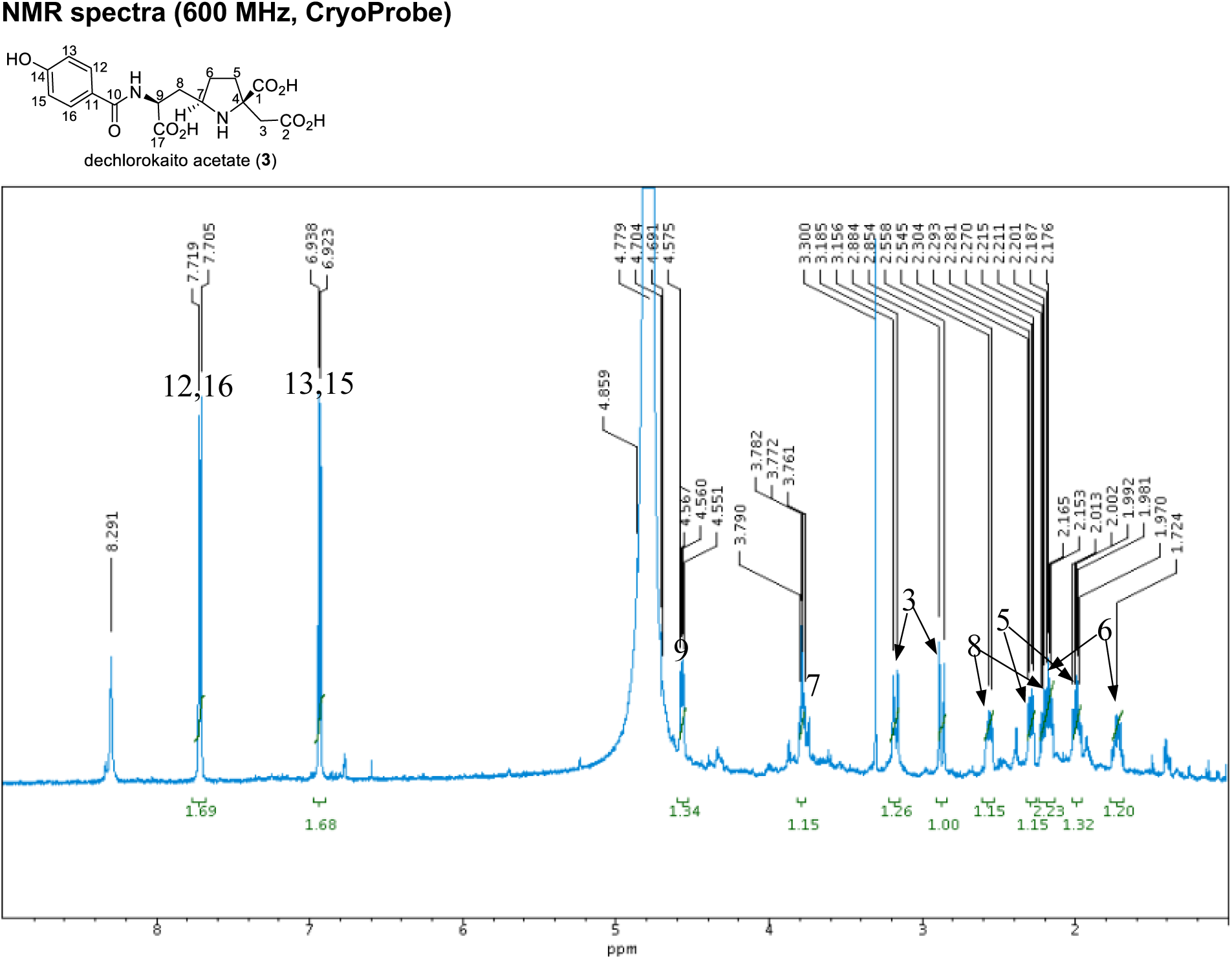
^1^H NMR spectrum of **3** in D_2_O

**Figure S7.**
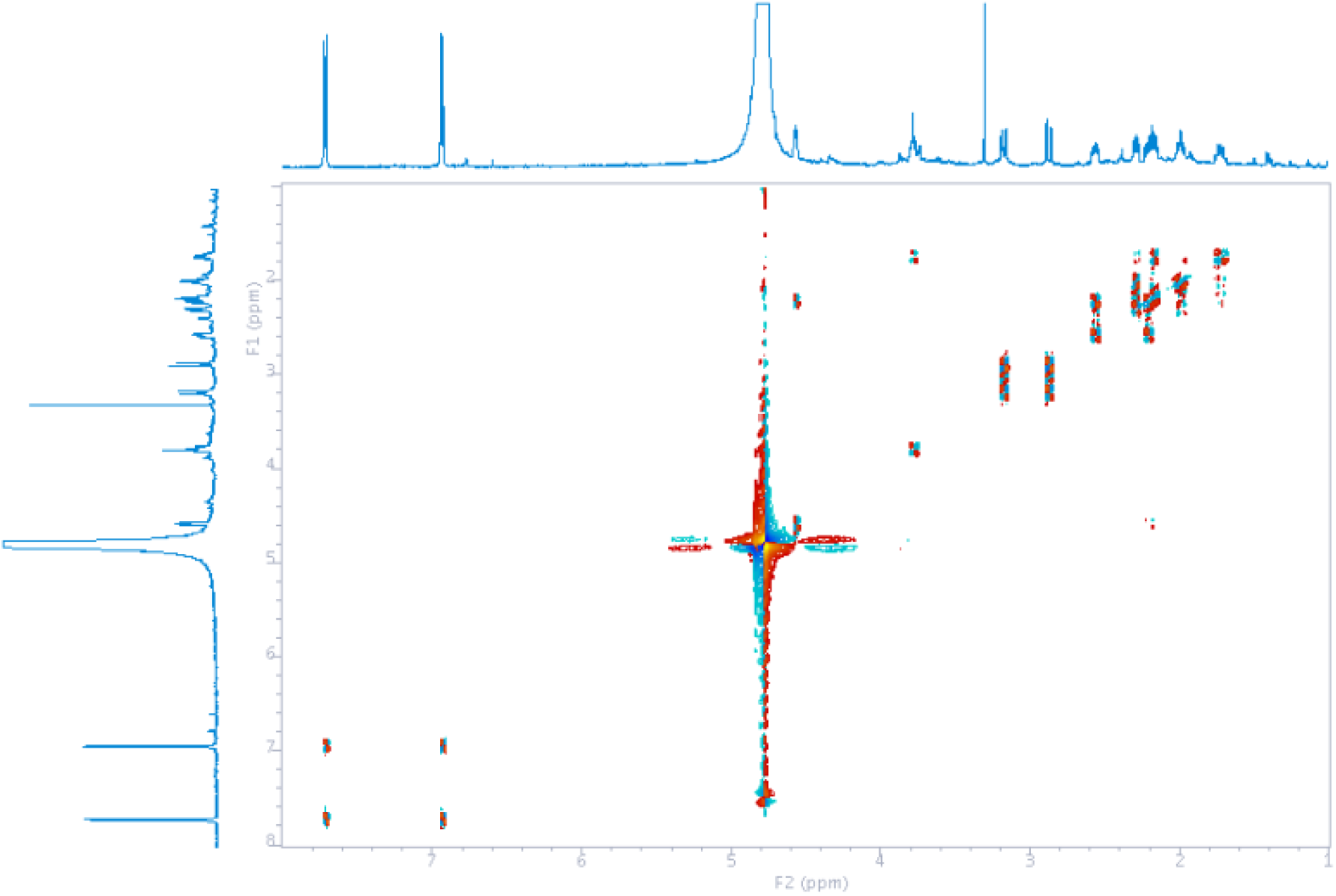
COSY spectrum of **3** in D_2_O.

**Figure S8.**
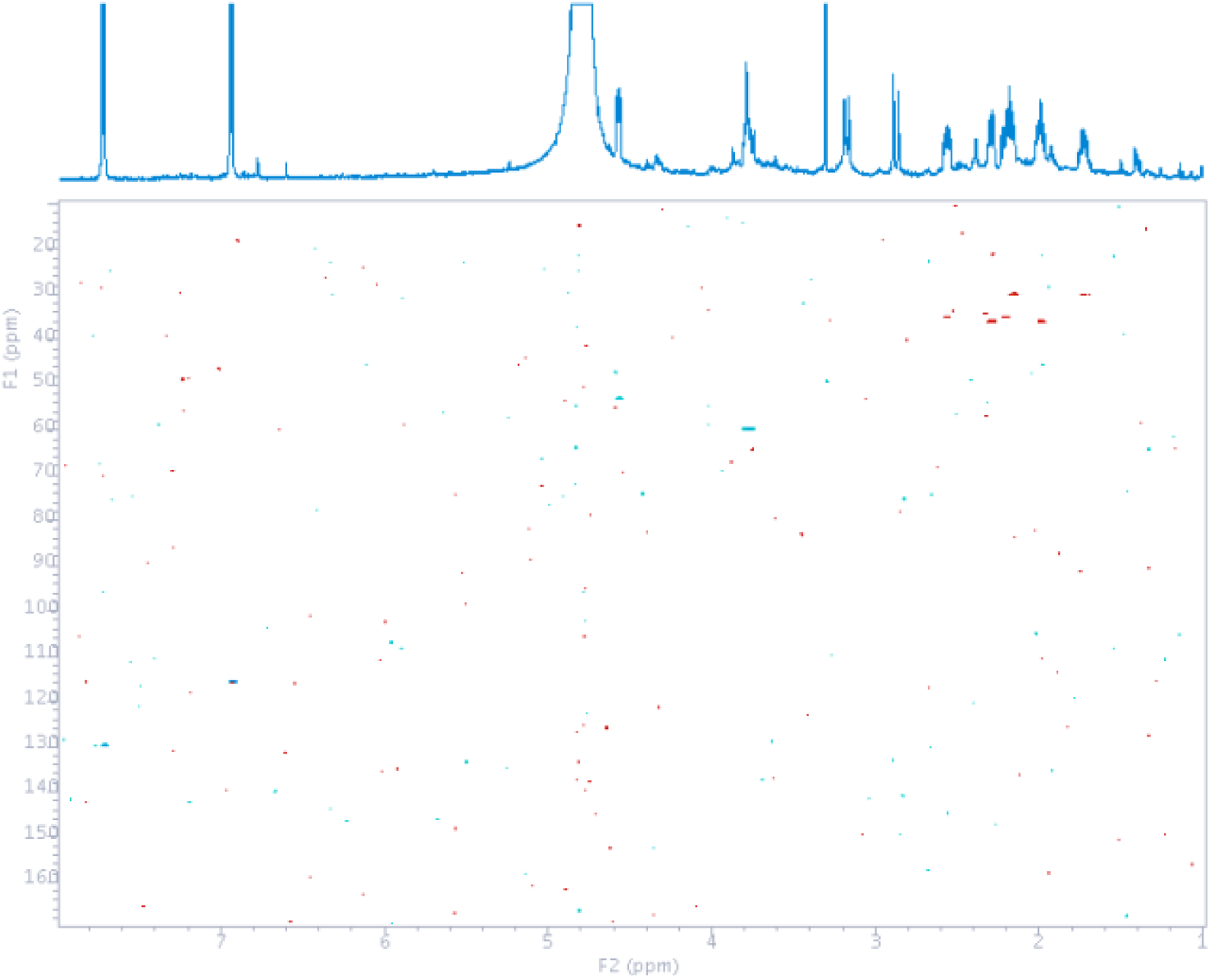
HSQC spectrum of **3** in D_2_O.

**Figure S9.**
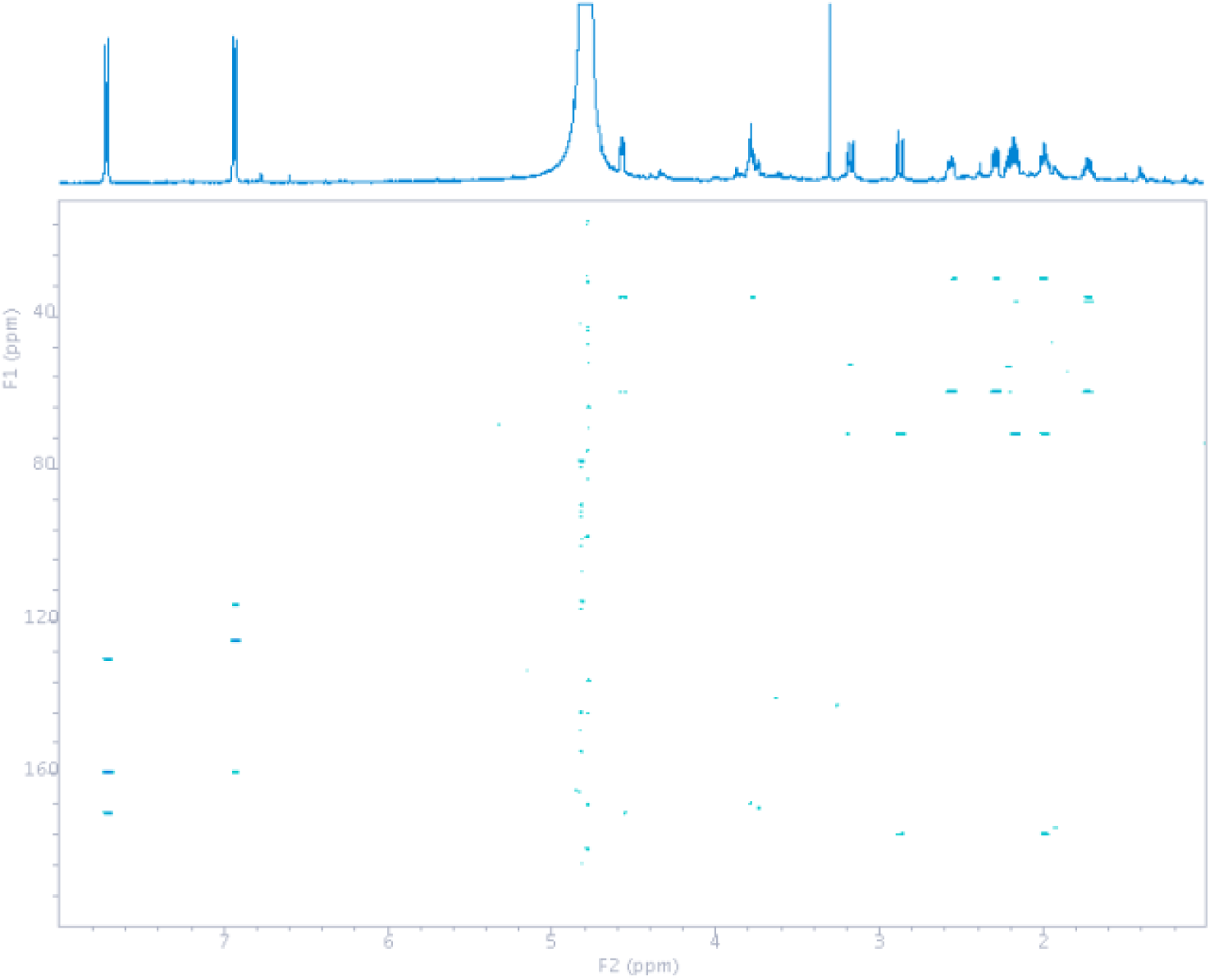
HMBC spectrum of **3** in D_2_O.

**Figure S10.**
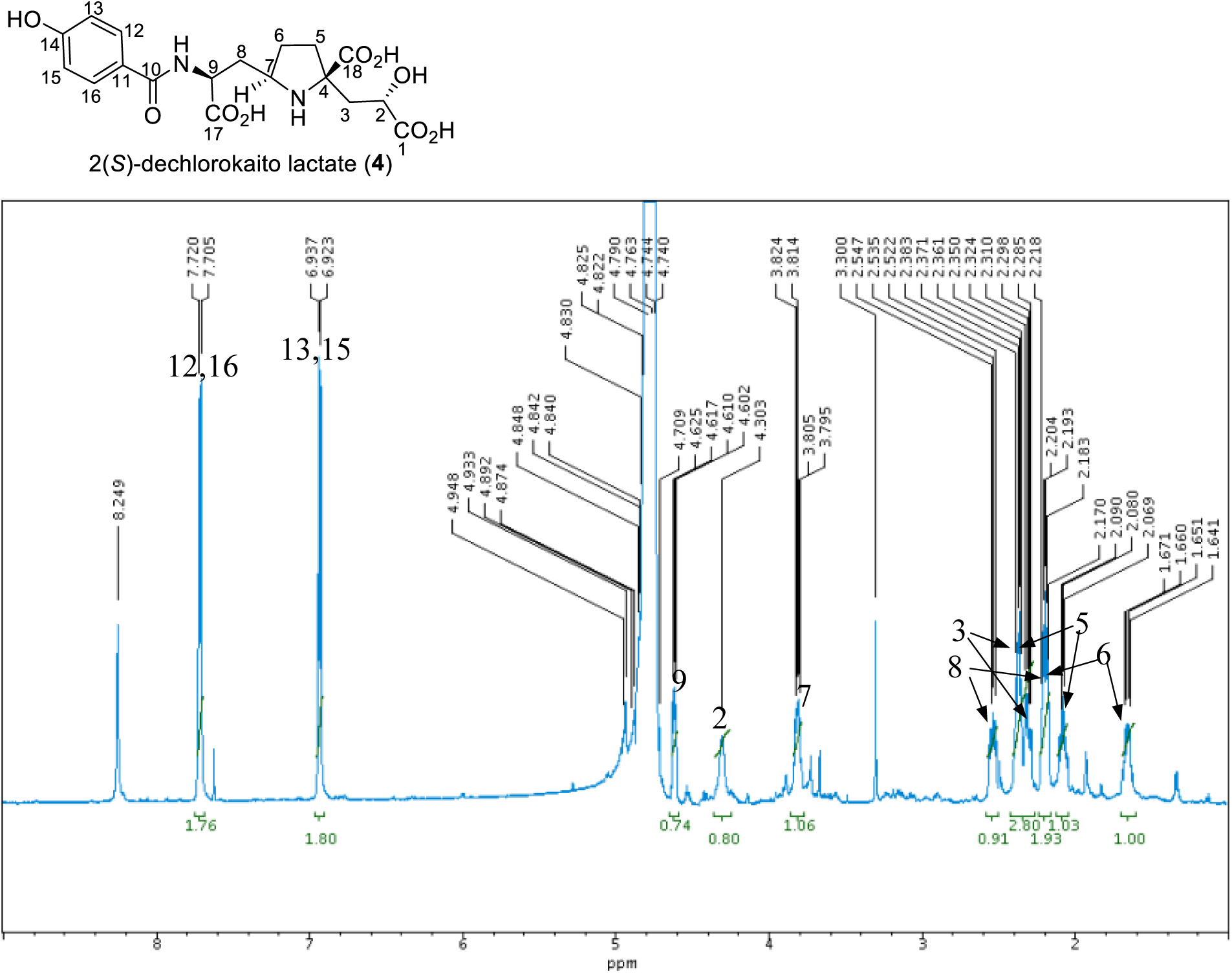
^1^H NMR spectrum of **4** in D_2_O

**Figure S11.**
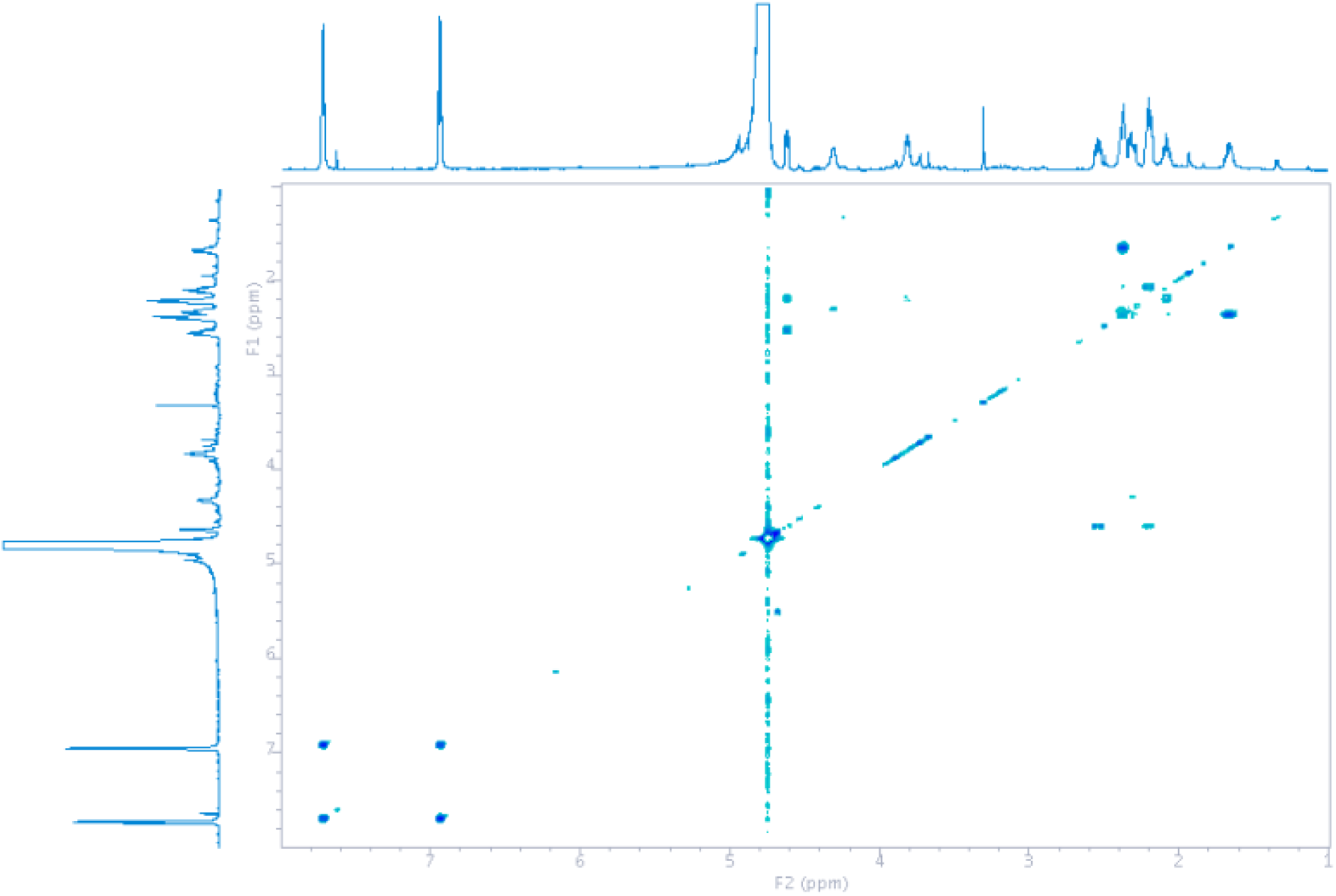
COSY spectrum of **4** in D_2_O.

**Figure S12.**
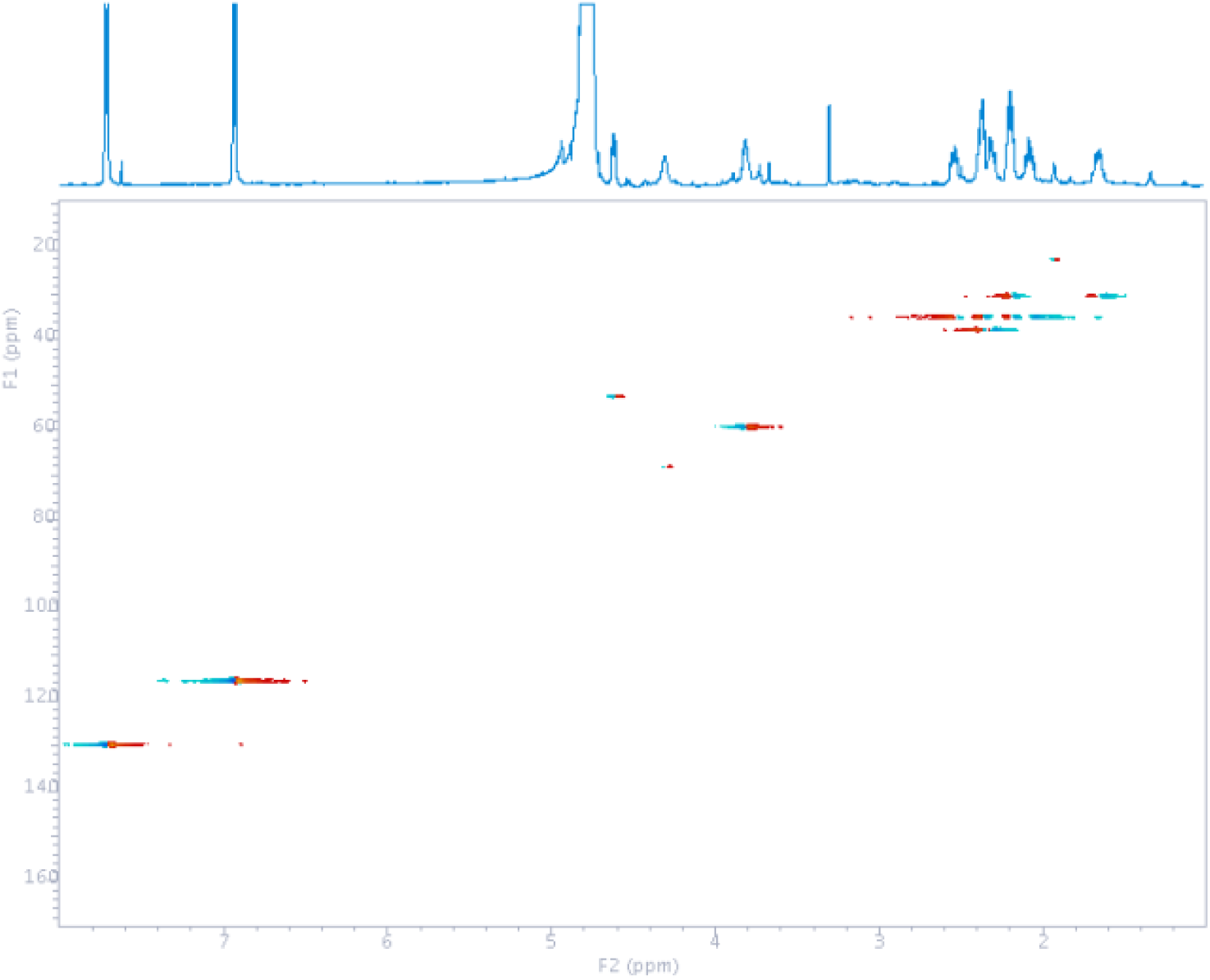
HSQC spectrum of **4** in D_2_O.

**Figure S13.**
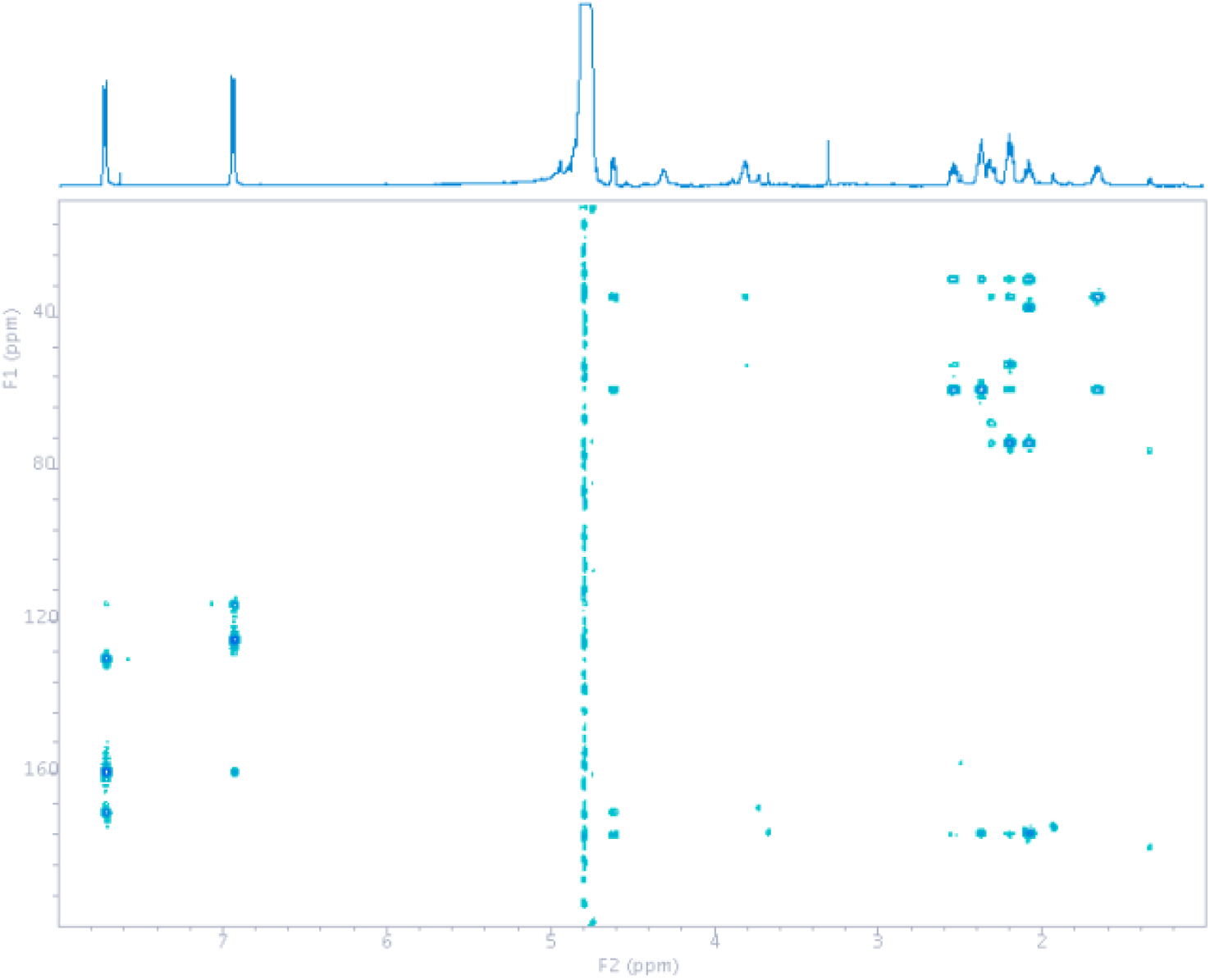
HMBC spectrum of **4** in D_2_O.

**Figure S14.**
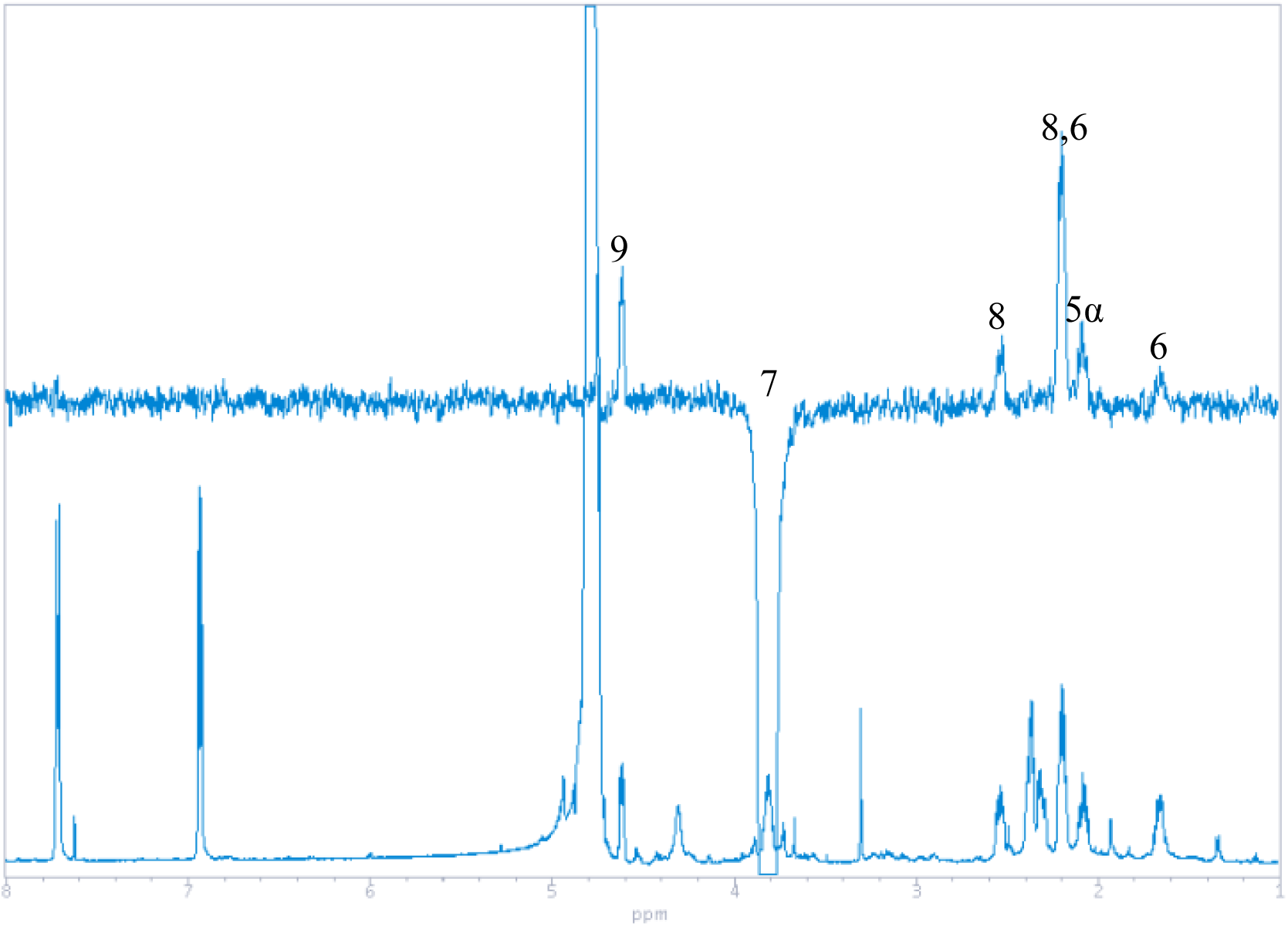
NOESY1D spectrum of **4** in D_2_O. Irradiated at 3.81 ppm (H-7).

**Figure S15.**
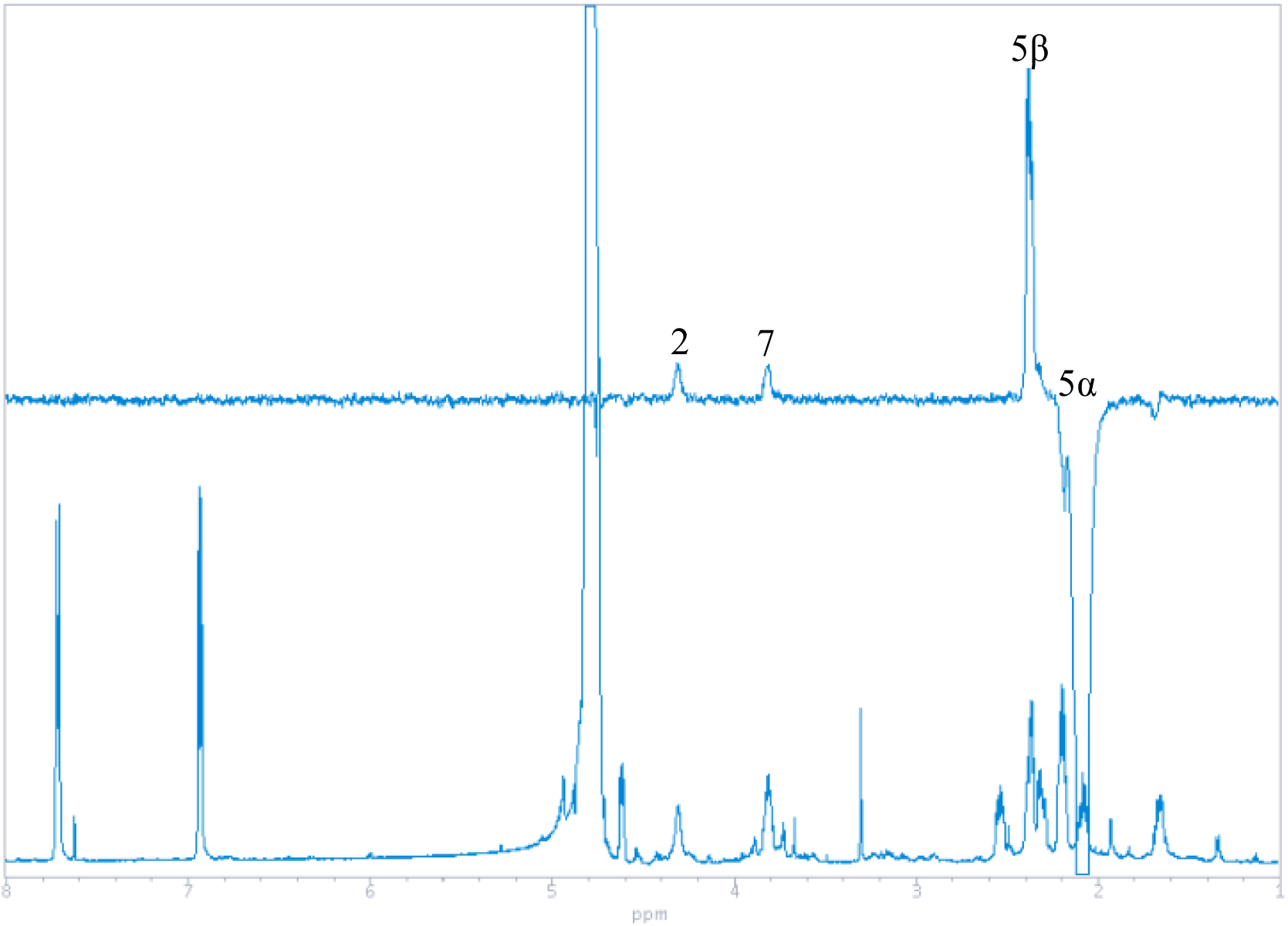
NOESY1D spectrum of **4** in D_2_O. Irradiated at 2.08 ppm (H-5α).

**Figure S16.**
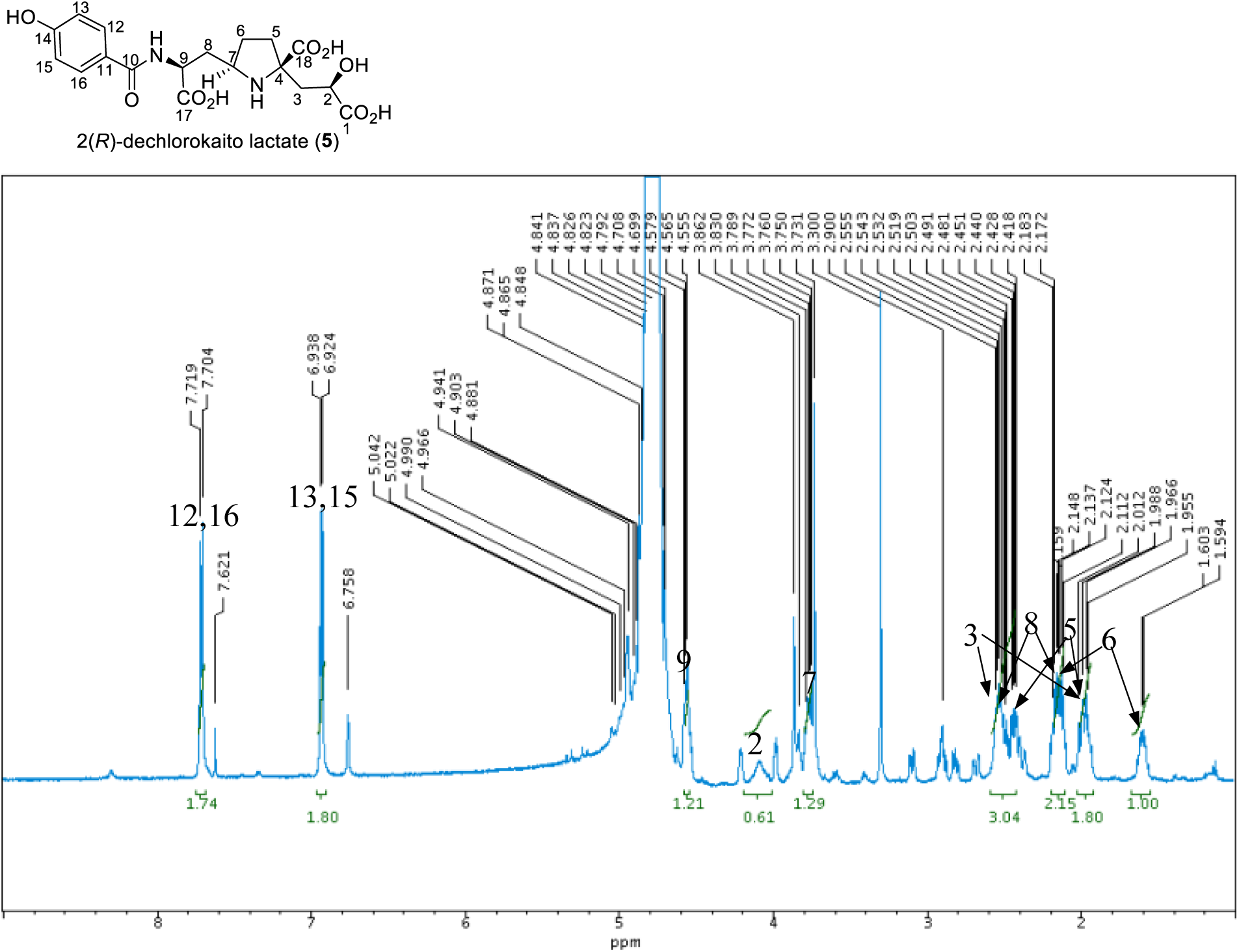
^1^H NMR spectrum of **5** in D_2_O

**Figure S17.**
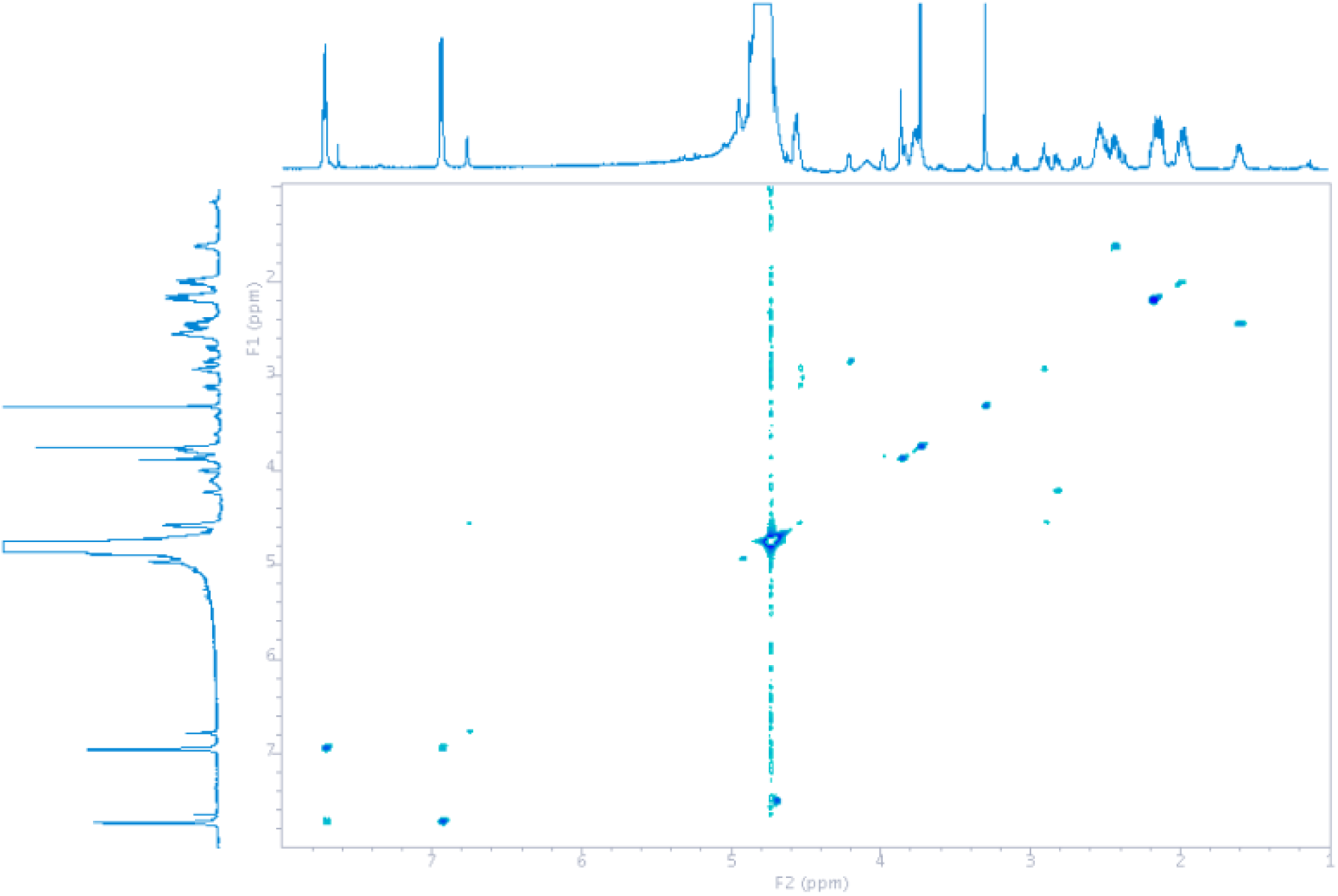
COSY spectrum of **5** in D_2_O.

**Figure S18.**
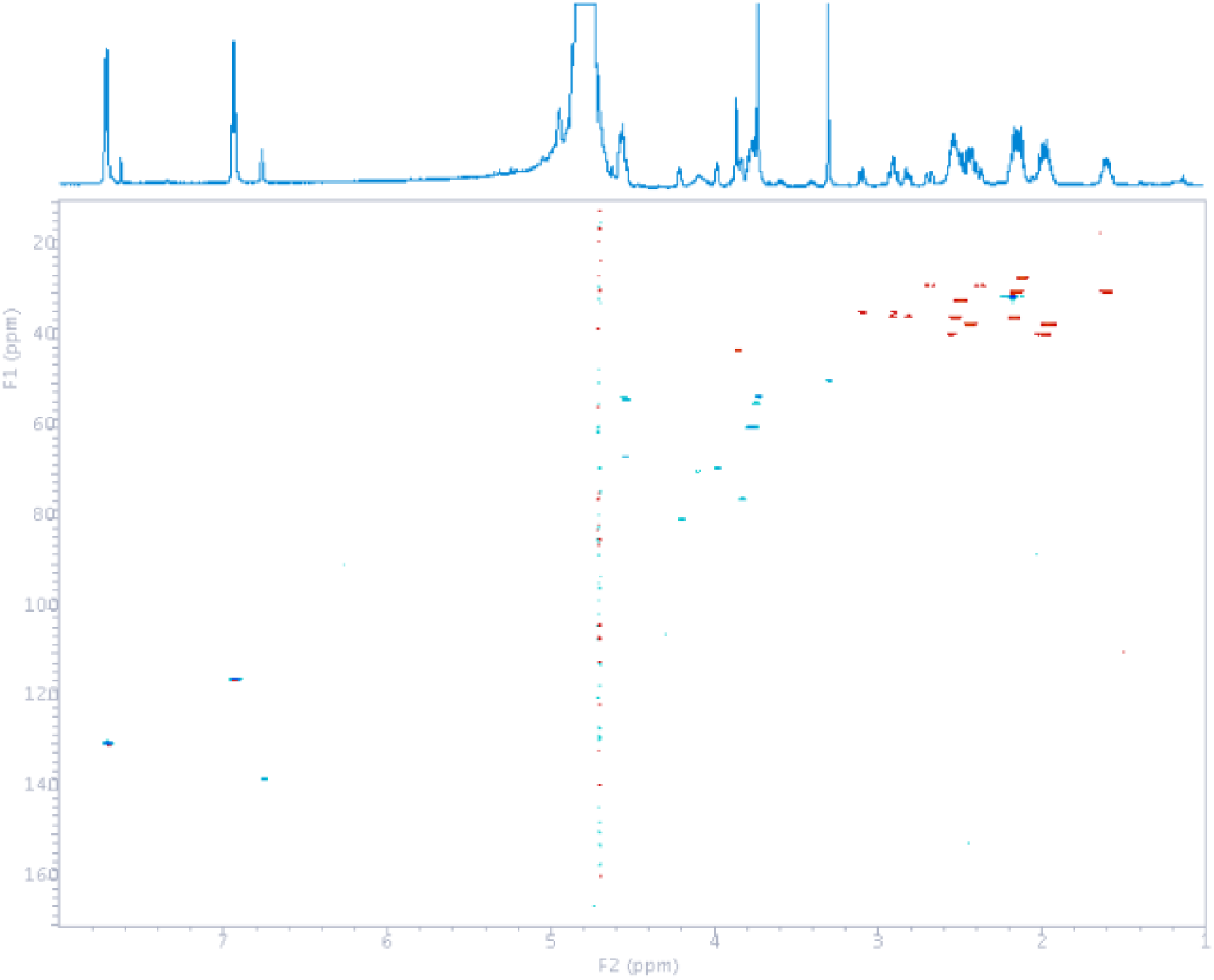
HSQC spectrum of **5** in D_2_O.

**Figure S19.**
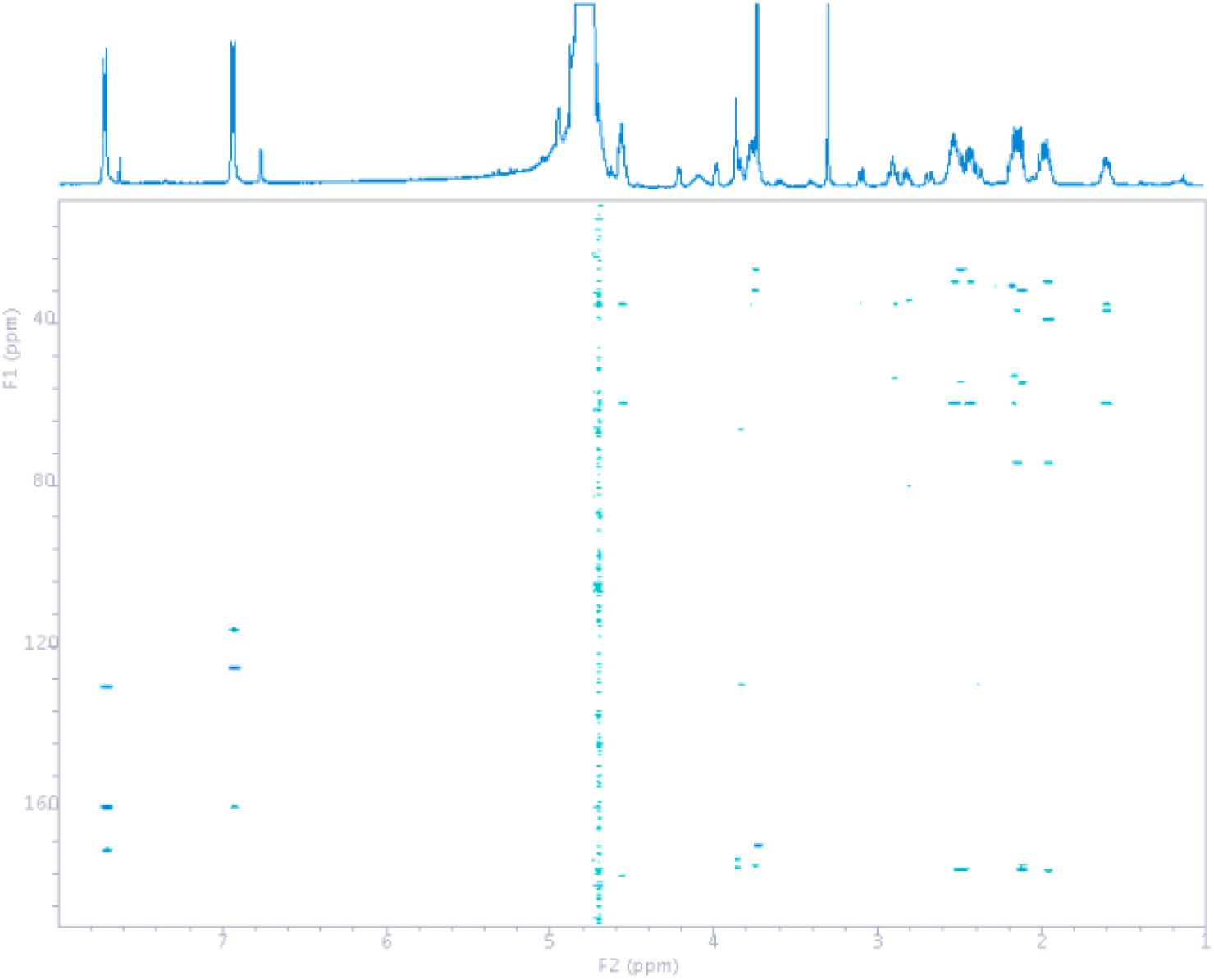
HMBC spectrum of **5** in D_2_O.

**Table S1.**
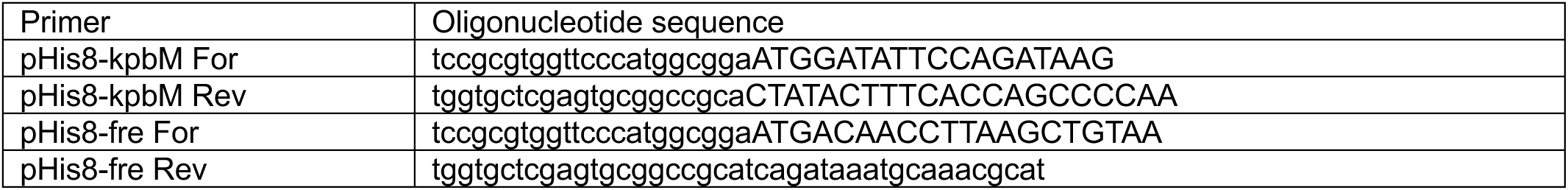
List of primers used in this study.

**Table S2.**
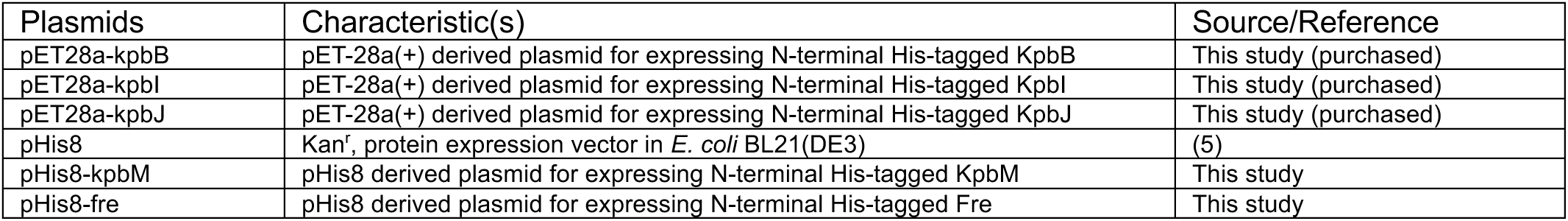
Plasmids used in this study.

**Table S3.** ^1^H-NMR data (*δ*_H_, *J* in Hz) of 3–5.

| position | **3** |  | **4** |  | **5** |  |
| --- | --- | --- | --- | --- | --- | --- |
| | $\delta_{\text{H}} J$ in Hz | | $\delta_{\text{H}} J$ in Hz | | $\delta_{\text{H}} J$ in Hz | |
| 2 | - |  | 4.3, brs |  | 4.1, brs |  |
| 3 | 2.87, d (18.2) | 3.17, d (17.6) | 2.31, m | 2.37, m | 2.01, m | 2.54, m |
| 4 | - |  | - |  | - |  |
| 5 | 1.98, m | 2.29, m | 2.08, m | 2.36, m | 1.96, | 2.43, m |
| 6 | 1.72, m | 2.17, m | 1.65, m | 2.19, m | 1.60, m | 2.12, m |
| 7 | 3.77, m |  | 3.81, m |  | 3.75, m |  |
| 8 | 2.21, m | 2.55, m | 2.20, m | 2.53, m | 2.17, m | 2.53, m |
| 9 | 4.56, dd (9.3, 4.7) |  | 4.61, dd (9.1, 5.0) |  | 4.56, m |  |
| 12, 16 | 7.71, d (8.5) |  | 7.71, d (8.5) |  | 7.71, d (8.8) |  |
| 13, 15 | 6.93, d (8.5) |  | 6.93, d (8.5) |  | 6.93, d (8.5) |  |

The signal of remaining CH<sub>3</sub>OD (3.30 ppm) was used as the internal reference.

**Table S4.** ^13^C-NMR data (*δ*_c_, type) of 3–5.

| position | 3 |  | 4 |  | 5 |  |
| --- | --- | --- | --- | --- | --- | --- |
| | $\delta_c$, type | HMBC | $\delta_c$, type | HMBC | $\delta_c$, type | HMBC |
| 1 | 175.6, C | 3, 5 | 178.5, C | 3 | N.D. | - |
| 2 | 174.2, C | 3 | 68.0, CH | 3 | 69.0, CH | 3 |
| 3 | 40.0, CH<sub>2</sub> |  | 37.5, CH<sub>2</sub> | 5 | 38.6, CH<sub>2</sub> | 5 |
| 4 | 70.2, C | 3, 5 | 72.6, C | 3, 5, 6 | 74.0, C | 5, 6 |
| 5 | 35.4, CH<sub>2</sub> | 6 | 34.7, CH<sub>2</sub> | 3, 6, 7 | 36.3, CH<sub>2</sub> | 6 |
| 6 | 29.5, CH<sub>2</sub> | 5, 8 | 30.0, CH<sub>2</sub> | 5, 7, 8 | 29.1, CH<sub>2</sub> | 5, 7, 8 |
| 7 | 59.2, CH | 6, 8 | 59.1, CH | 5, 6, 8, 9 | 59.3, CH | 5, 6, 8 |
| 8 | 34.4, CH<sub>2</sub> | 6, 7, 9 | 34.8, CH<sub>2</sub> | 6, 7, 9 | 34.8, CH<sub>2</sub> | 6, 7, 9 |
| 9 | 52.4, CH | 8 | 52.3, CH | 7, 8 | 52.6, CH | 8 |
| 10 | 170.0, C | 12, 16 | 170.2, C | 9, 12, 16 | 170.1, C | 9, 12, 16 |
| 11 | 125.0, C | 13, 15 | 124.9, C | 13, 15 | 124.8, C | 13, 15 |
| 12 | 129.4, CH | 16 | 129.7, CH | 16 | 129.5, CH | 16 |
| 13 | 115.3, CH | 15 | 115.6, CH | 12, 15 | 115.4, CH | 12, 15 |
| 14 | 159.3, C | 12, 13, 15, 16 | 159.6, C | 12, 13, 15, 16 | 159.4, C | 12, 13, 15, 16 |
| 15 | 115.3, CH | 13 | 115.6, CH | 13, 16 | 115.4, CH | 13, 16 |
| 16 | 129.4, CH | 12 | 129.7, CH | 12 | 129.5, CH | 12 |
| 17 | 176.4, C | 9 | 175.8, C | 8, 9 | 176.4, C | 8, 9 |
| 18 | - |  | 176.2, C | 3, 5 | 175.1, C | 5 |

N.D. = Not detected because of the limited amounts.
$\delta_c$ were determined based on HSQC and HMBC data. The signal of remaining $^{13}\text{CH}_3\text{OD}$ (49.0 ppm) was used as the internal reference.

